# ATaRVa: Analysis of Tandem Repeat Variation from Long-Read Sequencing data

**DOI:** 10.1101/2025.05.13.653434

**Authors:** Abishek Kumar Sivakumar, Anukrati Sharma, Sriram Sudarsanam, Sai Siddharth Krishnavajjhala, Harriet Dashnow, Akshay Kumar Avvaru, Divya Tej Sowpati

**Affiliations:** CSIR Centre for Cellular and Molecular Biology, Hyderabad, India; Academy of Scientific and Industrial Research, Ghaziabad, India; Department of Biomedical Informatics, University of Colorado Anschutz, Aurora, CO, USA

**Keywords:** Tandem repeats, microsatellites, genotyping, long-read sequencing, STRs, VNTRs, Pathogenicity

## Abstract

Tandem Repeats (TRs) are contiguous repetitions of DNA motifs whose variations modulate key cellular processes and traits. Their expansions are causally linked to over 70 neurological disorders in humans. Historically, genotyping TRs has been challenging due to their low sequence complexity and allele lengths exceeding typical short-read sequencing capabilities. While long-read sequencing improves TR genotyping by spanning large expansions and sequencing native DNA molecules, existing long-read genotypers often are inaccurate, computationally inefficient, and show platform-specific biases. Here, we present ATaRVa, a genotyper that achieves superior accuracy across both PacBio and Oxford Nanopore platforms, while running ∼15x faster than existing tools. Beyond genotyping, ATaRVa features sequence-level motif decomposition and methylation profiling. Evaluated across diverse datasets and genome-wide catalogs, ATaRVa especially excels in low coverage datasets, and accurately identifies known pathogenic expansions in clinical samples. Importantly, by quantifying uninterrupted target motifs in a population scale analysis, the tool systematically resolves false positives caused by benign, interrupted alleles. In addition, we present VisuaMiTRa, an auxiliary interface that enables interactive visualization of motif-level decomposition and base-level methylation of the TR alleles. By combining high computational efficiency with sequence-level structural awareness, ATaRVa provides a highly scalable framework for TR analysis in both population cohorts and clinical settings.

## Introduction

Tandem Repeats (TRs) are highly polymorphic genomic sequences composed of contiguous repetitions of DNA motifs. Based on the motif size, TRs can be categorised into Short Tandem Repeats (STRs, 1-6 bp) and Variable number tandem repeats (VNTRs, >= 7 bp) (Fan & Chu, 2007; J. M. Wright, 1994; Ziaei Jam et al., 2023). These regions can show high structural complexity, harboring multiple distinct motifs or a nested structure (Avvaru et al., 2025; Weisburd et al., 2026). TRs contribute to several key biological processes such as splicing, methylation, and transcription factor binding (Horton et al., 2023; S. E. Wright & Todd, 2023). Their inherently high mutation rates, ranging from 10^-5^ to 10^-3^ per locus per generation, lead to extensive polymorphisms in both their length and sequence composition. Following the discovery of CGG repeat expansion in *FMR1* gene associated with fragile X syndrome (FXS) (Verkerk et al., 1991), studies have found over 70 STR locations associated with Mendelian diseases such as ataxias, Huntington’s disease (HD), and frontotemporal dementia (Danzi, Xu, Fazal, Dolzhenko, Pellerin, Weisburd, Reuter, et al., 2025; Depienne & Mandel, 2021; Gall-Duncan et al., 2022). Pathogenic expansions can span several kilobases and drive disease via distinct mechanisms like loss of function, gain of function or alternate splicing. Notably, STR expansions classically referred to as ‘full mutations’ cause disease by inducing CpG hypermethylation and gene silencing (Gall-Duncan et al., 2022; Malik et al., 2021).

Although repeat lengths exceeding pathogenic thresholds typically exhibit full penetrance, their clinical severity varies widely. Recent studies suggest that sequence composition, in particular the presence of non-canonical motifs or interruptions, can alter the effect of expanded allele and determine the disease penetrance (Huang et al., 2022; Latham et al., 2014; G. E. B. Wright et al., 2019). In HD, while age at disease onset correlates with the size of pure CAG repeats, mouse models show that CAA interruptions within CAG repeats reduce the somatic expansion and modulate disease risk and severity. This variability in pathogenicity due to interruptions is also seen in fragile X syndrome (FXS), where AGG interruptions stabilize the repeat (Yrigollen et al., 2012). Conversely, sequence variations can trigger pathogenicity as well; in familial adult myoclonic epilepsy (FAME), large expansions of pure TTTTA motifs remain benign, but presence of other motifs such as TTTGA and TTTCA is pathogenic (Urabe et al., 2025).

Historically, tandem repeats, particularly VNTRs, have been difficult to genotype using short-read sequencing (SRS) data, as these reads often lack sufficient non-repetitive flanks to achieve precise mapping to the reference (Javadzadeh et al. 2025). The advent of long-read sequencing (LRS) technologies from Pacific Biosciences (PacBio) and Oxford Nanopore Technologies (ONT) has solved this physical limitation, producing reads spanning tens of kilobases that encapsulate entire TR loci. Leveraging this benefit, several long-read TR genotypers have recently emerged to identify and profile these previously inaccessible regions (Chiu et al. 2021; Dolzhenko et al. 2024; Ziaei Jam et al. 2024; Lougheed et al. 2026).

However, translating these technological advances into population-scale discoveries is currently hindered by bottlenecks spanning computational efficiency, genotyping accuracy, and sequence resolution. At the biobank scale, existing tools suffer from prohibitive processing times, high memory (RAM) consumption, and architectures optimized for specific sequencing platforms. Furthermore, many tools exhibit biases in phasing and determining zygosity, introducing systematic errors and compromising genotyping accuracy (Aliyev et al., 2026). More importantly, while current tools report copy numbers and allele sequences, they fail to resolve the complex internal sequence composition of massively expanded alleles across diverse populations (Morato Torres et al., 2022). For example, TRGT reports the copy numbers only of predefined motifs (Dolzhenko et al., 2024), TREAT reports information only for a single representative motif (Tesi et al., 2024), and *vamos* relies on static regex heuristics tied to a predefined reference motif (Ren et al., 2023). These targeted approaches inherently disregard novel, non-canonical motifs and random sequence interruptions. This limitation is particularly critical for TRs embedded in highly polymorphic “Variation Clusters” (VCs), including eight disease-associated loci (*ATXN8OS*, *FGF14*, *NOP56*, *CNBP*, *HTT*, *BEAN1*, *LRP12*, and *EIF4A3*), where internal sequence composition, rather than the allele length alone, determines disease severity (Aston & Dion, 2025; Weisburd et al., 2026). Establishing a framework that captures precise motif composition independent of existing knowledge is essential to reveal how internal sequence structures contribute to population-level traits.

Here, we present ATaRVa (Analysis of Tandem Repeat Variation), a sequencing technology-agnostic TR genotyping and analysis framework. ATaRVa implements a highly optimized, read-wise processing architecture that drastically improves processing speed and reduces memory consumption across both PacBio and ONT platforms. By combining repeat-aware flank-realignment with unbiased haplotype clustering, ATaRVa achieves a superior genotyping accuracy across platforms, zygosities, sequencing depth, and in complex regions. Beyond deriving accurate allele-lengths, ATaRVa decomposes TR alleles into precise motif-level representation and reports base-level CpG methylation to resolve differential epigenetic states. Furthermore, to facilitate immediate clinical and biological interpretation, we introduce an auxiliary web application, VisuaMiTRa, which directly utilizes ATaRVa outputs to visually resolve sequence composition and differential methylation profiles across cohorts.

## Results

### Algorithm Architecture and Core Features

ATaRVa optimizes genotyping by leveraging the high density of genome-wide TR catalogs, and processes each read only once to record information for all the TR loci it covers. It takes a long-read alignment file, a tandem repeats coordinate file, and the reference genome fasta file as input. ATaRVa iterates through high-quality reads from target chromosomal chunks, and records information for each covered TR locus, including flanks, insertion/deletion (InDel) positions, base modification data, and single-nucleotide variants (SNVs).

ATaRVa maintains three primary data structures, for read-, locus-, and SNV-specific information (Fig 1). A TR locus is genotyped once all supporting reads have been collected, dropping locus-specific information post-genotyping to maintain a low memory footprint. Genotyping begins with local realignment of the flanking regions using a SIMD optimised Smith–Waterman algorithm (Zhao et al., 2013) to identify and incorporate misaligned repeat insertions. Supporting reads are subsequently phased using nearby informative (heterozygous) SNVs (Fig S1a). If informative SNVs are absent, reads are clustered using HDBSCAN (*HDBSCAN*, n.d.) based on the read repeat length and edit distance to the reference repeat (Fig S1b). Consensus sequences for each haplotype are generated using abPOA (Gao et al., 2021). Finally, the consensus alleles undergo motif decomposition (see Motif Decomposition section), and the average CpG methylation probability is calculated across all haplotype reads (Fig S2a).

**Figure 1:**
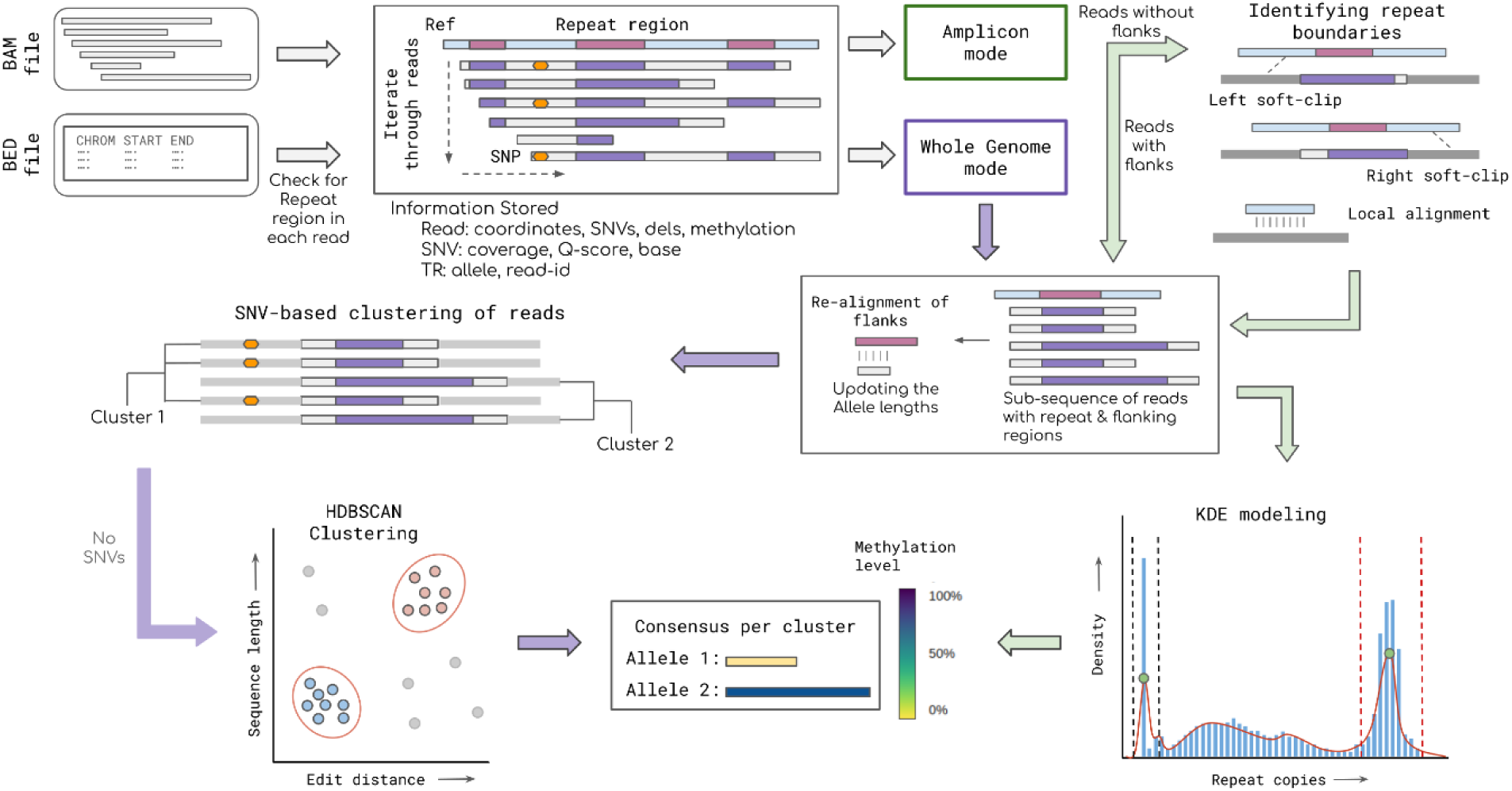
Overview of ATaRVa algorithm. The flowchart highlights the main steps of the algorithm, including the two genotyping modes of ATaRVa - Whole Genome, and Amplicon. A detailed description of the algorithm and various steps is provided in the Supplementary Information.

The Amplicon mode tackles target-amplified long-read sequencing of highly unstable pathogenic TR loci. This mode employs a locus-wise processing strategy. Crucially, to recover massive repeat expansions that exceed standard read alignments, this mode actively searches within soft-clipped read segments for locus-flanking sequences (Fig S2b). Given the typical lack of informative SNVs in targeted sequencing, the Amplicon mode estimates genotypes by modeling read repeat copy numbers using Kernel Density Estimation (KDE) (*KernelDensity*, n.d.). Using a custom scoring scheme, the two most significant KDE peaks are retained as candidate alleles. If only one significant peak is identified then the dominant cluster undergoes a check for subclusters using HDBSCAN analysis. When multiple subclusters are present, the two largest are selected as representative alleles; otherwise, the locus is considered homozygous.

By providing motif-level decomposition (`DS` tag) and allele-specific methylation profiling (`MA` and `MV` tags) in the output VCF, ATaRVa extends TR genotyping beyond simple allele length estimation, enabling the detailed characterization of sequence interruptions alongside structural and epigenetic variations. To support the exploration and interpretation of these patterns, we developed an auxiliary tool, VisuaMiTRa (Visualization of Motifs & Methylation in Tandem Repeat Alleles; Fig S3a and S3b). VisuaMiTRa directly parses the ATaRVa-generated VCF to extract motif decomposition, methylation data, and locus metadata. This encoded information is transformed into an expanded visual representation of motif boundaries, sequence interruptions, and overlaid 5mC methylation profiles. In addition, the interface provides customization options that allow users to modify color palettes, legends, and font settings. VisuaMiTRA supports both single-sample and multi-sample analyses, enabling users to seamlessly interpret TR sequence and epigenetic variations at population scale. A detailed description of the ATaRVa algorithm and its underlying parameters is provided in the Supplementary Information.

### Genotyping Accuracy Across Sequencing Platforms

We evaluated the accuracy of ATaRVa on three distinct 30x whole-genome sequencing (WGS) datasets of the widely used Genome-In-A-Bottle (‘Genome in a Bottle’, 2012) HG002 sample: High Fidelity reads from Pacific Biosciences (PacBio HiFi), Oxford Nanopore Technologies (ONT) Duplex, and ONT Simplex. The evaluated tandem repeat catalog comprised 5,599,658 TR loci from TRExplorer v0.2 (Weisburd et al., 2026). We compared ATaRVa against four established long-read base genotypers: LongTR, Medaka Tandem, STRkit and TRGT (PacBio only). All tools showed a mean genotyping rate of ∼99%, with the exception of STRkit (96.6%; Table S1).

We compared tool-derived allele lengths against the ground-truth allele lengths derived from the high-quality phased assembly of the HG002 sample from the T2T consortium Q100 project (Hansen et al., 2025). For this, we selected a subset of 5,184,662 loci exhibiting exactly two alleles in the reference assembly. On this subset, all tools maintained a genotyping rate of ∼100%, except for STRkit (Table S1). On the PacBio HiFi dataset, ATaRVa achieved an exact-match concordance rate of 86.4%, significantly outperforming TRGT (82%), STRkit (74%) and LongTR (67%; Fig 2a). When allowing for a ±1 bp tolerance, the concordance of ATaRVa improved to 97.5%, maintaining its lead over LongTR (93.8%), TRGT (91.4%), and STRkit (86.1%).

**Figure 2:**
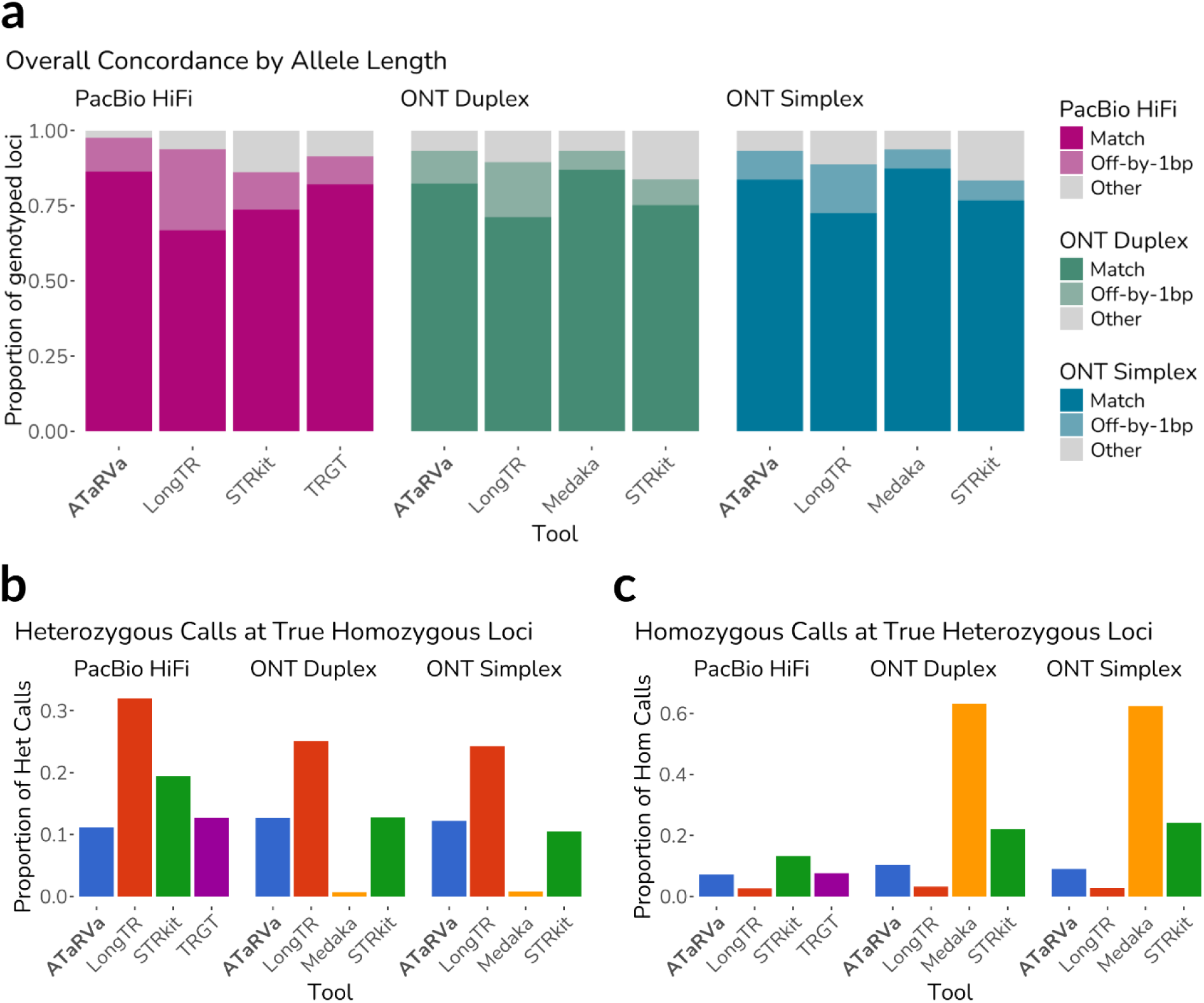
Genotyping accuracy and concordance of ATaRVa compared to existing long-read genotypers on the HG002 genome. **a)** Overall genotyping performance across three 30× sequencing datasets: PacBio HiFi, ONT Duplex, and ONT Simplex. Stacked bars represent the proportion of genotyped loci classified as an exact match to the assembly-derived ground truth (darkest shade), within a ±1 bp tolerance (medium shade), or other errors (light grey). Note that TRGT and Medaka Tandem were evaluated exclusively on PacBio HiFi and ONT datasets, respectively. **b)** The proportion of false heterozygous calls generated at true homozygous loci, highlighting the bias of LongTR toward heterozygous calls across all tested technologies. **c)** The proportion of false homozygous calls generated at true heterozygous loci, revealing Medaka’s extreme bias toward homozygous calls on ONT datasets. In contrast, ATaRVa maintains balanced, platform-agnostic accuracy across all zygosity states.

ATaRVa demonstrated a similar robust performance on ONT datasets. For ONT Simplex data, ATaRVa and Medaka Tandem achieved the highest concordance rate of 93% (±1 bp tolerance), outperforming LongTR (89%) and STRkit (83.4%; Fig 2a). This ranking hierarchy was consistently maintained across the ONT Duplex data. Global concordance results for all tools across the three datasets are summarized in Table S1.

### Resolving Genotyping Biases by Zygosity

To investigate the effect of TR allele zygosity on genotyping accuracy, we stratified tool performance across 4,613,712 homozygous loci and 570,950 heterozygous loci, as defined by the HG002 assembly.

For homozygous loci in the PacBio HiFi dataset, ATaRVa achieved the highest ±1 bp concordance (98.9%), outperforming LongTR (96.3%), TRGT (94.7%), and STRkit (88.7%; Fig S4a). On ONT Simplex data, Medaka Tandem led with 98.1% concordance, followed by ATaRVa (95.6%), LongTR (91.6%), and STRkit (86.3%; Fig S4a). Notably, Medaka Tandem also achieved the highest exact-match concordance (95.4%) for these loci, exceeding ATaRVa by 8.4 percentage points, a trend also seen in the ONT Duplex data (Table S2).

However, performance rankings shifted substantially for heterozygous loci. On PacBio HiFi data, ATaRVa maintained the highest concordance rate (86.0%), exceeding the next best tool by 11.8 percentage points (LongTR; 74.2%), followed by TRGT (65.2%), and STRkit (63.6%). In contrast to its strong performance at homozygous loci, Medaka Tandem showed the lowest concordance for heterozygous loci on ONT Simplex data (57.6%). ATaRVa again achieved the highest Simplex heterozygous concordance (73.8%), followed by LongTR (66.4%) and STRkit (58.0%), with Duplex data mirroring this pattern (Table S3 and Fig S4b).

To understand these divergent accuracies, we examined the proportion of incorrect genotype calls for each zygosity class (Fig 2b and 2c). For homozygous loci, LongTR exhibited the highest mean error rate (27.1%), followed by STRkit (14.2%), TRGT (12.6%; PacBio only), and ATaRVa (12.0%), while Medaka Tandem demonstrated a negligible error rate (0.7%; ONT only). However, this trend inverted completely for heterozygous loci: Medaka Tandem showed the highest mean error rate (62.7%), followed by STRkit (19.8%), ATaRVa (8.9%), TRGT (7.6%), and LongTR (2.8%; Table S4).

These results reveal distinct algorithmic biases: LongTR is strongly biased toward heterozygous calls, consistent with previous reports (Aliyev et al., 2026). Conversely, Medaka Tandem exhibits a severe bias toward homozygous calls, heavily inflating its apparent global concordance. Overall, ATaRVa and TRGT displayed the most balanced, unbiased performance across both heterozygous and homozygous loci, maintaining error rates near 10%.

### Performance Across Challenging Loci and Sequencing Depths

Given that homopolymers comprise approximately one-third of the TR catalog, we evaluated genotyping accuracy specifically on this subset of ∼1.6 million loci (Fig S5a). For PacBio HiFi, ATaRVa achieved the highest ±1 bp concordance (97.9%), followed by LongTR (94.3%), TRGT (93.5%), and STRkit (62.4%). For ONT data, Medaka Tandem achieved the highest concordance (92.0%), followed by ATaRVa (86.0%), LongTR (82.3%), and STRkit (55.1%; Table S5). This superior performance of Medaka Tandem is likely attributable to its specialized neural network–based error correction model. Exclusion of homopolymer loci improved exact-match concordance and reduced off-by-one errors for all tools except LongTR, which retained a relatively high off-by-one error rate (∼8% in HiFi, ∼3% for ONT), further confirming its heterozygous-calling bias (Table S6 and Fig S5b).

We next assessed performance across varying allele lengths (Fig 3a). Across platforms, genotyping accuracy predictably declined as allele length increased. However, ATaRVa demonstrated the most stable performance across all categories. In heterozygous loci across both HiFi and ONT datasets, ATaRVa consistently achieved the highest concordance, with TRGT showing the steepest decline in accuracy for alleles >1000 bp. In homozygous loci, ATaRVa led the HiFi benchmarks, while Medaka Tandem led the ONT benchmarks for longer alleles, though this is heavily influenced by aforementioned homozygous bias of Medaka. A similar trend was observed when we stratified our analysis by repeat motif length (Fig S6). ATaRVa consistently ranked highest for heterozygous loci across both platforms. Predictably, all tools struggled with homopolymers and extremely large motifs (>500 bp), but ATaRVa maintained greater stability in these edge cases. Conversely, TRGT exhibited the largest performance decline for large motifs.

**Figure 3:**
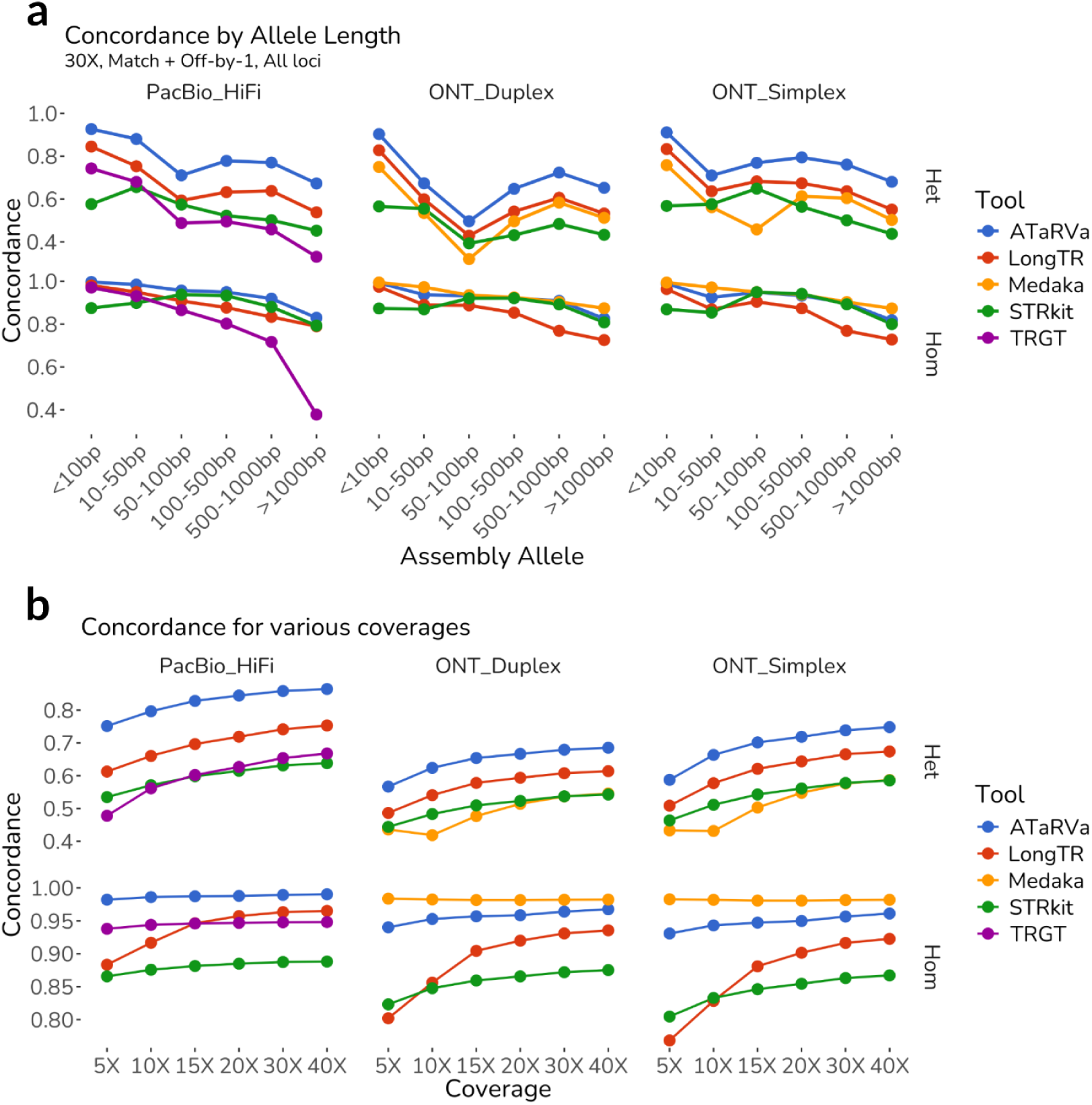
Genotyping accuracy across allele lengths and sequencing depths. **a)** Overall genotyping concordance (calculated as exact match + ±1 bp tolerance) stratified by reference assembly allele length and locus zygosity. Top row: heterozygous loci; bottom row: homozygous loci. Concordance trajectories demonstrate ATaRVa’s (blue) sustained accuracy across allele length categories, particularly highlighting its superior performance across heterozygous loci regardless of the underlying sequencing platform. **b)** Impact of sequencing coverage (5× to 40×) on overall concordance (exact match + ±1 bp tolerance). Performance is stratified by sequencing platform (PacBio HiFi, ONT Duplex, ONT Simplex) and locus zygosity. Top row: heterozygous; bottom row: homozygous. ATaRVa consistently outperforms existing algorithms across platforms and zygosity states, demonstrating a distinct advantage at low sequencing depths (5X–15X), particularly for heterozygous loci.

Finally, to understand the effect of sequencing coverage, we evaluated the tools on subsampled datasets ranging from 5x to 40x (Fig 3b). ATaRVa achieved the highest concordance across all coverage levels for both homozygous and heterozygous HiFi data, effectively resolving loci even at 5× depth. The accuracy of LongTR improved sharply from 5x to 15x coverage, followed by a more gradual increase at higher coverages. However, it consistently remained below ATaRVa. In ONT datasets, Medaka Tandem dominated homozygous calls across all depths but consistently failed at heterozygous loci. These results indicate that ATaRVa is the most accurate and balanced tool across sequencing depths.

### Multi-Sample Validation and Mendelian Consistency

To confirm these metrics generalize beyond the de facto HG002 benchmark, we evaluated ATaRVa on PacBio HiFi datasets from five additional Human Pangenome Reference Consortium (HPRC) samples (HG00438, HG00621, HG00673, HG02622, and HG02630). Consistent with the HG002 results, ATaRVa outperformed LongTR, TRGT, and STRkit across all length concordance thresholds, allele lengths, and zygosity classes (Fig 4a and Fig S7).

**Figure 4:**
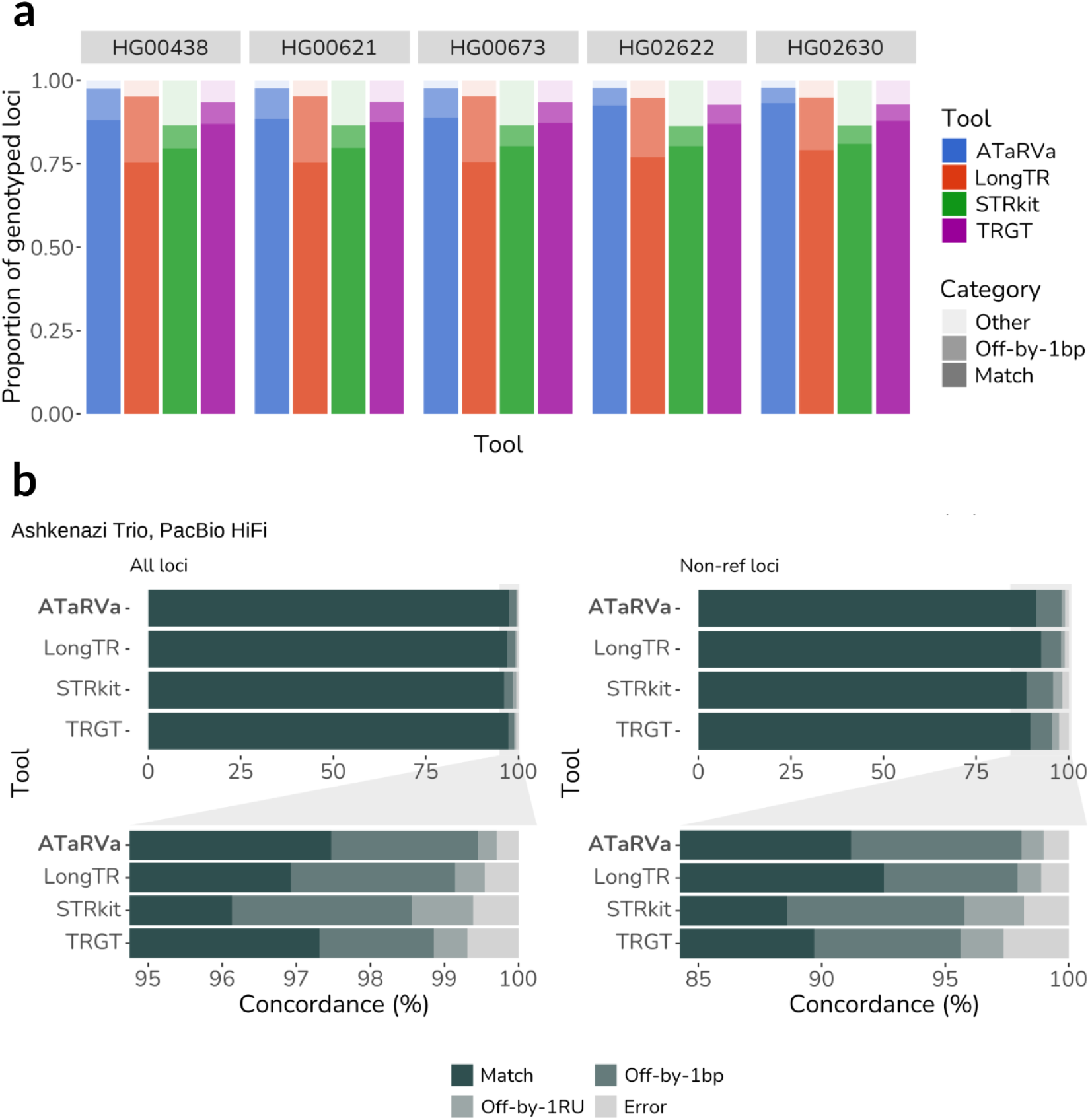
Multi-sample validation and Mendelian consistency. **a)** Multi-sample validation showing the proportion of accurately genotyped loci across five diverse HPRC genomes (PacBio HiFi data). Stacked bars display exact matches (darkest shade), calls within a ±1 bp tolerance (medium shade), and other discrepancies (lightest shade). **b)** Mendelian inheritance concordance evaluated within the Ashkenazi Jewish trio (HG002, HG003, HG004) using PacBio HiFi data. Performance is assessed across all loci (left) and restricted to non-reference loci (right) to eliminate reference-bias. The bottom charts provide a magnified view of the upper concordance percentiles. Alleles are categorized as an exact match, off-by-1 bp, off-by-1 repeat unit (RU; equivalent to one motif difference), or error (>1 RU difference).

We further assessed Mendelian concordance using the PacBio HiFi data of the Ashkenazi Jewish trio (HG002, HG003, and HG004). Medaka Tandem was excluded in this analysis due to platform incompatibility. Alleles were classified as exact match, off-by-1 bp, off-by-1 motif, or error (>1 motif difference). Across all loci, ATaRVa achieved a Mendelian concordance of 97.47% for exact matches and 99.45% for ±1 bp tolerance between parent and child alleles (Fig 4b and Table S7), outperforming the other tools.

When restricting the analysis strictly to loci that were non-reference (i.e., excluding loci homozygous for the reference allele in all three family members), concordance expectedly decreased. However, ATaRVa remained the most accurate, achieving 91.18% exact-match and 98.08% ±1 bp concordance (Fig 4b and Table S7). In this non-reference subset, LongTR reported approximately 700,000 more loci than TRGT, aligning with its known bias to over-call heterozygous genotypes. Crucially, ATaRVa consistently exhibited the lowest proportion of alleles in the ‘error’ category across both the complete and non-reference subsets, indicating a superior ability to correctly resolve complex repeat structures and track allelic inheritance without introducing systematic length artifacts.

### Detection and High-Resolution Profiling of Pathogenic Expansions

To evaluate the ability of ATaRVa to detect pathogenic repeat expansions, we analyzed 33 samples carrying known pathogenic expansions across 11 disease-associated genes (*AR, ATN1, ATXN1, ATXN3, DMPK, FMR1, FXN, C9ORF72, PABPN1, RFC1,* and *HTT*). These datasets were obtained from multiple sources (PacBio website; Dolzhenko et al., 2024; Stevanovski et al., 2022) (see Data Availability). Of the 33 samples, 29 were generated using targeted PacBio sequencing, 2 were PacBio whole-genome sequencing (WGS) datasets, and 2 were targeted Oxford Nanopore Technologies (ONT) adaptive sampling sequencing datasets. All five genotyping tools were executed across the cohort (using their “targeted sequencing” preset where available; see Methods) to assess their sensitivity in identifying expanded alleles (Fig 5a).

**Figure 5:**
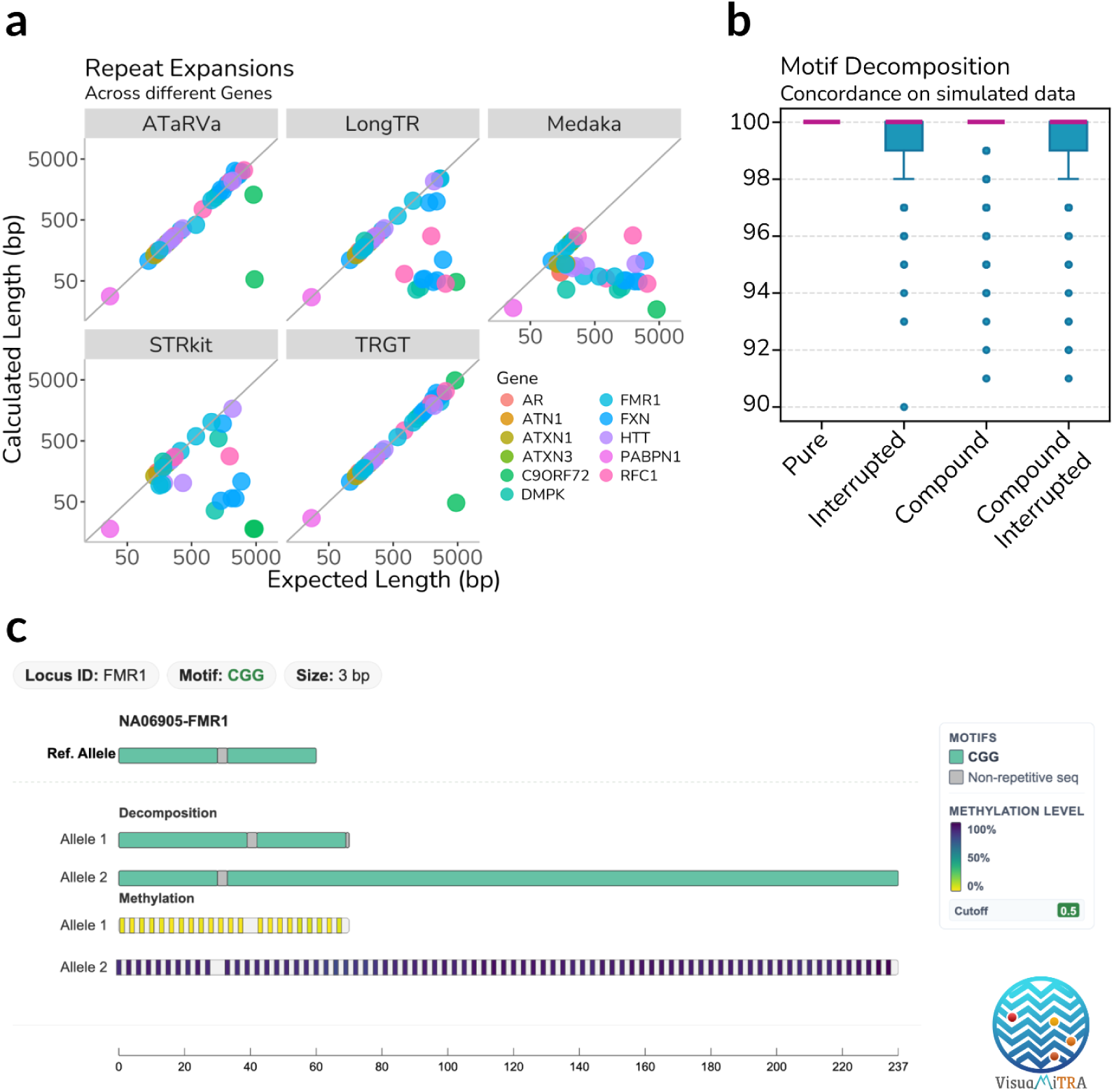
High-resolution profiling of pathogenic expansions. **a)** Comparison of calculated versus expected allele lengths across samples with known expansions at disease-associated loci. ATaRVa accurately resolves massive expansions, matching the diagnostic sensitivity of TRGT. Conversely, LongTR, Medaka, and STRkit systematically underestimate allele sizes or fail entirely at extended lengths (>500 bp). Axes are log-scaled. **b)** Concordance of ATaRVa’s sequence-level motif decomposition evaluated on simulated data. High accuracy is maintained across diverse and complex repeat architectures, including pure, interrupted, compound, and compound-interrupted alleles. **c)** Interactive visualization of an *FMR1* expansion using the VisuaMiTRa interface. The platform aligns the motif-level structural architecture (top tracks, denoting the expanded CGG motif) directly with base-resolution DNA methylation profiling (bottom tracks), facilitating rapid clinical interpretation of both sequence composition and epigenetic state.

Utilizing its specialized `--amplicon` mode, designed to recover massively expanded alleles via unaligned soft-clip processing and robust artifact handling, ATaRVa correctly identified diagnostic categorizations for 32 of the 33 samples. Of these, ATaRVa identifies the same allele length as the original studies in 30 samples. In two additional cases (one *FMR1* and one *C9ORF72*), ATaRVa underestimated the absolute allele length; however, the reported sizes remained within their established pathogenic thresholds (Table S8). TRGT correctly identified the expansions in 32 samples. However, it must be noted that for 23 of the 33 evaluated samples, specifically those sourced from PacBio reference datasets, the ground-truth expansion status was originally established using TRGT itself. Despite this inherent methodological alignment between the dataset curation and TRGT’s performance, ATaRVa demonstrated equivalent pathogenic sensitivity, failing to detect an expansion in only a single sample. Notably, the single expansion missed by ATaRVa was also missed by TRGT: the ∼4.7 kb *C9ORF72* expansion in sample ND11494. This expansion completely eluded all evaluated tools because it was supported only by a single spanning read. Both LongTR and STRkit consistently underestimated the size of larger pathogenic expansions or failed to resolve them entirely. Furthermore, although Medaka Tandem is primarily optimized for ONT data, its licensing does not explicitly restrict its use to ONT sequencing. In this cohort (where 31 out of 33 samples are PacBio HiFi data), Medaka Tandem failed to accurately identify the vast majority of pathogenic expansions, systematically misreporting them as short alleles.

From a clinical standpoint, ATaRVa offers a diagnostic advantage by resolving not only total allele length but also the underlying allelic sequence architecture. ATaRVa offers motif-level decomposition of the consensus TR allele sequence. We have demonstrated motif decomposition performance on the simulated data generated for four different categories (Methods; Fig S8a and S8b). Benchmarking on 1,000 simulated sequences per category, the script achieved a 100% concordance rate for the pure repeat dataset (Fig 5b). Furthermore, it accurately resolved and decomposed more than 98% of the simulated sequences across the remaining three complex categories (Interrupted, Compound, and Compound-Interrupted TRs).

To demonstrate the utility of this fine-scale resolution, we examined the *HTT* locus in sample NA13509 (Fig S8c). The motif architectures of the two alleles were resolved as follows:

● Allele 1: (CAG)_15_-CAACA-(GCC)_2_-ACC-(GCC)_9_-G
● Allele 2: (CAG)_73_-CAACA-(GCC)_2_-ACC-(GCC)_6_-G

This sequence-level resolution allows users to immediately differentiate the stable wild-type allele (CAG₁₅) from the pathogenic expanded allele (CAG₇₃), while simultaneously tracking the arrangement and copy number of adjacent interrupting motifs. Capturing this internal architecture facilitates the investigation of repeat interruptions, which are increasingly recognized as critical modifiers of repeat instability and disease severity.

Beyond structural decomposition, ATaRVa integrates DNA methylation profiling, which can be interactively explored via VisuaMiTRa. We utilized VisuaMiTRa to examine the *FMR1* CGG repeat expansion in sample NA06905 (Fig 5c). The interface clearly delineated distinct methylation patterns between the two alleles. The expanded allele (237 bp) exhibited extensive CpG hypermethylation with a mean methylation level of 91%. In stark contrast, the normal-length allele showed a mean methylation level of only 3%, with minimal methylation observed across its CpG sites. This allele-resolved visualization highlights the direct structural association between repeat expansion and localized hypermethylation, perfectly mirroring the known pathogenic mechanism of Fragile X syndrome, wherein *FMR1* expansions trigger promoter hypermethylation and subsequent transcriptional silencing (Verkerk et al., 1991).

### Computational Efficiency, Scalability, and Execution Overhead

To ensure an equitable baseline comparison given the highly variable multithreading support across tools, processing speed was evaluated in single-threaded mode on a representative subset of 100,000 TR loci (Fig 6a). On 30× PacBio HiFi data, ATaRVa processed approximately 435 loci/s, outperforming the next fastest tool, TRGT (∼392 loci/s), by 10%. The performance advantage was even more pronounced on ONT data. On 30× ONT Simplex data, ATaRVa maintained a speed of ∼294 loci/s, more than 16-fold faster than Medaka Tandem (∼17 loci/s) and over an order of magnitude faster than the remaining tools.

**Figure 6:**
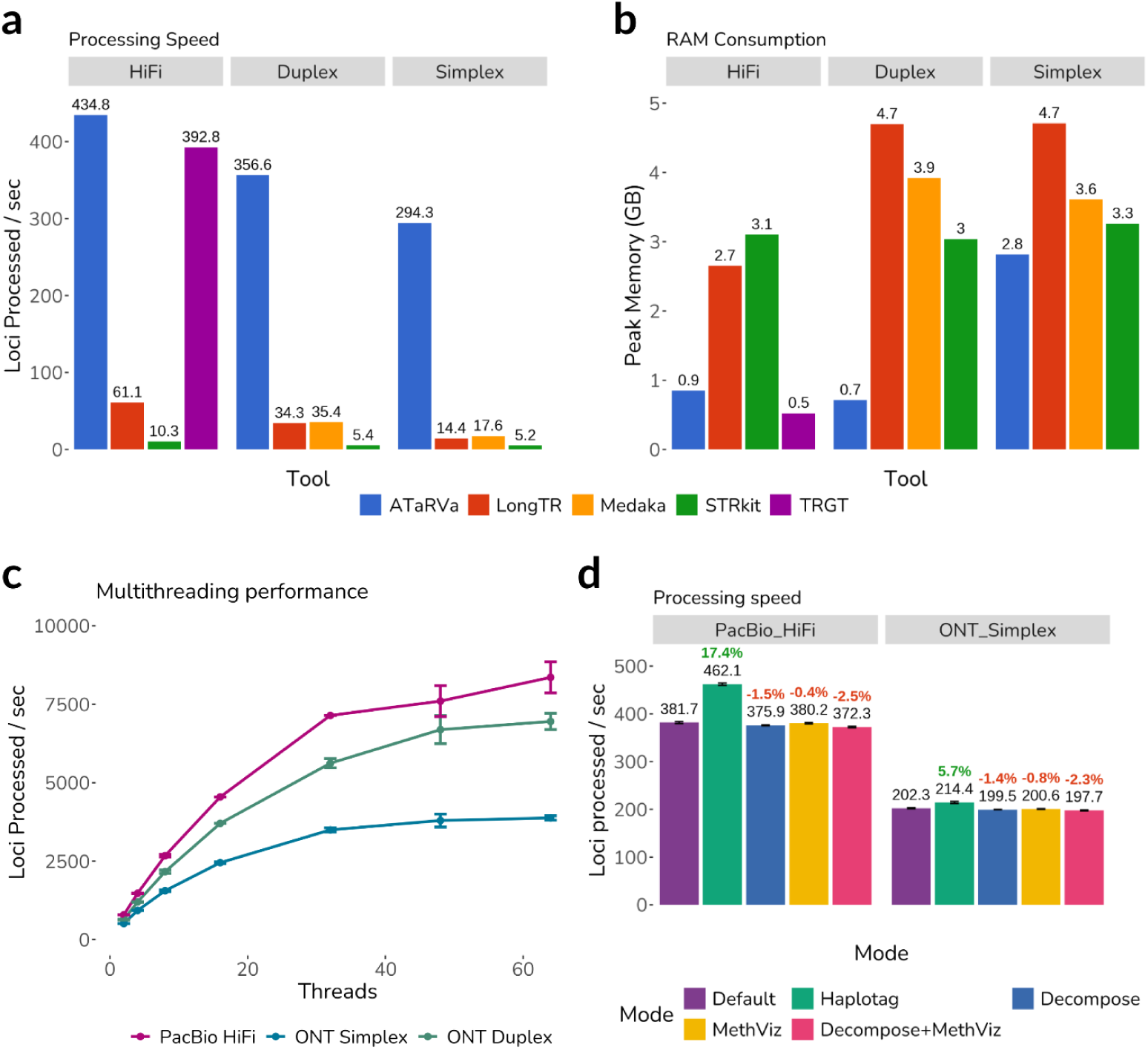
Computational efficiency and scalability of the ATaRVa framework. **a)** Processing speed, measured as loci processed per second, across PacBio HiFi, ONT Duplex, and ONT Simplex platforms. ATaRVa operates up to ∼15 times faster than cross-platform competitor tools on ONT datasets, while matching the high-speed performance of TRGT on PacBio HiFi. **b)** Peak memory (RAM) consumption in gigabytes (GB). ATaRVa maintains a minimal memory footprint across all technologies, utilizing a fraction of the RAM required by LongTR, Medaka, and STRkit. **c)** Multithreading performance across varying thread counts. The framework scales efficiently up to 32 threads across all tested sequencing technologies. **d)** Impact of advanced analytical modules on execution time. Activating additional profiling features, including sequence-level decomposition (Decompose) and DNA methylation analysis (MethViz), incurs negligible performance penalties (<3% reduction in speed) compared to the default execution mode, while providing phased (Haplotag) inputs slightly improves processing speed.

The relative reduction in ATaRVa’s processing speed and the concurrent increase in peak memory consumption on ONT Simplex data (2.8 GB) compared to PacBio HiFi (0.9 GB) and ONT Duplex data (0.7 GB) are attributable to the raw sequencing error profile of Simplex reads. Elevated error rates generate a massive influx of apparent single-nucleotide variants (SNVs). Because ATaRVa dynamically extracts and evaluates these SNVs during its phasing and clustering steps, the computational and memory overhead inherently scales with base-calling noise.

Despite this error-induced overhead, ATaRVa remained highly memory-efficient. On PacBio HiFi data, TRGT required the least peak memory (0.5 GB), closely followed by ATaRVa (0.9 GB; Fig 6b). Across all ONT datasets, ATaRVa exhibited the lowest peak memory utilization of any evaluated tool. Specifically on ONT Simplex data, ATaRVa consumed 13% less memory than STRkit, which maintained a static ∼3.0 GB footprint across all modalities, and over 20% less memory than both Medaka Tandem (3.6 GB) and LongTR (4.7 GB). We further assessed ATaRVa’s scalability via multithreading. Execution time scaled efficiently with increasing core counts up to 32 CPUs; beyond this threshold, speed gains became marginal (Fig 6c), indicating that maximum throughput becomes constrained by hardware I/O and memory-access bandwidth rather than algorithmic limits.

To quantify the computational penalty of advanced repeat profiling, ATaRVa was executed with `--decompose`, `--methviz`, and a combined `--decompose --methviz` configuration. On both PacBio and ONT datasets, these auxiliary modules introduced a negligible runtime penalty of ∼2%, demonstrating that ATaRVa can concurrently resolve complex motif structures and epigenetic states without compromising its core speed advantage (Fig 6d).

Finally, we evaluated the impact of the `--haplotag` mode, which bypasses internal SNV extraction in favor of pre-phased haplotype tags. This optimization increased processing speed by 17% and 5% for PacBio HiFi and ONT data, respectively (Fig 6d). When evaluated against the full genome-wide catalog, `--haplotag` execution on PacBio HiFi data increased exact-match concordance by ∼3% without altering overall ±1 bp concordance (Fig S9a). Conversely, ONT Simplex data exhibited a modest ∼2% reduction in overall accuracy when relying on external haplotype tags, particularly for longer allele and motif lengths (Fig S9b and S9c). This divergence highlights the robustness of ATaRVa’s default internal SNV-based phasing strategy when navigating error-prone long-read data.

### Population-Scale Pathogenic TR Profiling of the 1000 Genomes Long-Read Cohort

To demonstrate the utility of ATaRVa for population-scale, sequence-level TR analysis, we genotyped 66 autosomal pathogenic TR loci (catalogued in the STRchive database) across 498 presumed healthy individuals from the 1000 Genomes Project Long-Read Sequencing Cohort (1KGP-LRSC) Gibson et al., 2026; Gustafson et al., 2024) sequenced with ONT. Genotyping was performed with `--decompose` and `--methviz` flags. ATaRVa output VCFs from 498 samples were merged using the “`atarva merge”` utility (Fig S10). Initially, using length alone, alleles were classified as benign, intermediate, or pathogenic based on established copy-number thresholds defined by STRchive (Hiatt et al., 2025) for the target motifs of the corresponding loci (Aston & Dion, 2025; Rajan-Babu et al., 2024). Consistent with the cohort’s healthy status, allele length distributions for the vast majority of loci fell within expected benign ranges. However, preliminary length-based screening flagged individuals with allele lengths in established pathogenic ranges across ten loci, corresponding to the genes *RFC1, BEAN1, SAMD12, HTT, XYLT1, DAB1, RAI1, FGF14, STARD7* and *TCF4* (Fig S11 and Table S9). Crucially, the pathogenicity of these loci is determined not solely by total allele length, but by the repeat sequence composition, such as the presence of pathogenic motifs and sequence interruptions. For example, *RFC1*, *BEAN1*, and *RAI1* each harbor a benign reference motif alongside one or more distinct pathogenic variant motifs causally linked to disease.

To overcome the limitations of simple length-based classification, we leveraged sequence-level resolution offered by ATaRVa to detect repeat interruptions and novel motifs. Furthermore, previous studies have shown that across highly variable TR loci, tracking the longest uninterrupted motifs than measuring the overall length provides more biologically meaningful comparison (Danzi, Xu, Fazal, Dolzhenko, Pellerin, Weisburd, Van De Vondel, et al., 2025). Here we introduce an analytical metric, the Longest Pure Motif (LPM), which quantifies the maximum uninterrupted tract of a specific target motif within an allele. The LPM metric, reported directly in the FORMAT field of the output VCFs, enables users to definitively separate true pathogenic expansions from heavily interrupted, biologically benign long alleles. We performed a detailed, sequence-level re-evaluation of the flagged alleles utilizing LPM and the decomposed sequences extracted from the DS tag.

### Resolving False Positives at the XYLT1 Locus

The promoter region of *XYLT1* harbors a GCC repeat; expansions exceeding 72 copies are causally associated with Baratela-Scott Syndrome (BSS) due to hypermethylation-induced gene silencing (LaCroix et al., 2019). However, the expansion is contained within an 238 bp sequence that is absent from the hg38 reference genome. Therefore, using length alone to assess pathogenicity includes the flanking non-GCC sequence and results in false positives. By extracting the longest uninterrupted GCC repeat segment using the LPM tag, we re-evaluated the initial length-based pathogenic classifications. The LPM segments revealed that none of the alleles initially flagged as pathogenic actually crossed the copy-number threshold for pure GCC expansions (Fig 7a and Fig S12). This demonstrates ATaRVa’s capacity to immediately resolve false-positive calls generated by traditional length-only estimation.

**Figure 7:**
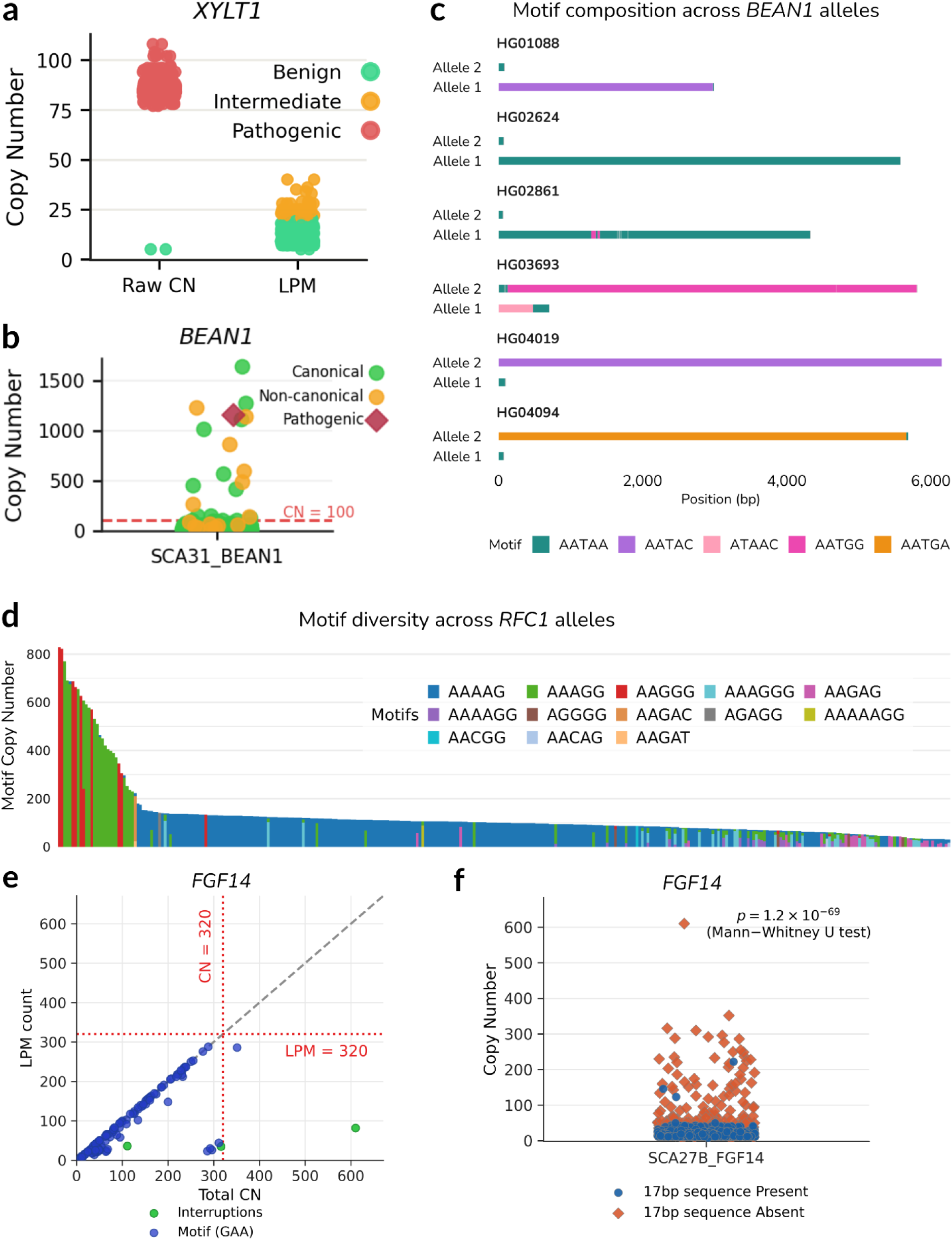
Sequence-level resolution of tandem repeat expansions and structural false positives. **a)** Reclassification of *XYLT1* expansions. Relying strictly on raw copy number (Raw CN) inaccurately classifies numerous alleles as pathogenic. Applying the Longest Pure Motif (LPM) metric reveals extensive internal interruptions, correctly reclassifying these alleles into benign and intermediate categories. **b)** Copy number distribution at the SCA31 (*BEAN1*) locus, distinguishing between alleles harboring canonical, non-canonical (previously unreported), and pathogenic expanded motifs. The red dashed line denotes the pathogenic threshold (CN = 100). **c)** Motif decomposition of massive *BEAN1* expansions across six representative individuals. The structural profiles demonstrate that extreme expansions at this locus are frequently driven by highly pure, distinct non-canonical motifs (e.g., AATAC, ATAAC). **d)** Motif diversity and structural complexity across expanded *RFC1* alleles, illustrating extensive sequence heterogeneity within the population cohort. **e)** Total copy number versus LPM count at the *FGF14* locus. Several alleles exceed the total length pathogenic threshold (CN = 320, red dashed lines) but are resolved as structurally benign when evaluated by their LPM, representing structural false positives. **f)** Distribution of *FGF14* copy number stratified by the presence or absence of an upstream protective 17-bp sequence, with statistical significance determined by a two-sided Mann-Whitney U test.

### Exploring Motif Diversity at the BEAN1 Locus

Expansions of non-reference AATGG or AATAG motifs beyond 110 copies at the *BEAN1* locus are associated with Spinocerebellar Ataxia Type 31 (SCA31) (Ishikawa 2023). We reassessed the pathogenicity of the flagged length-based expansions at this locus by integrating motif decomposition and LPM information. As expected, the majority of alleles in the cohort carried the benign AATAA reference motif (Fig 7b). Notably, we identified a single truly expanded allele harboring the pathogenic AATAG motif. Additionally, a subset of alleles were found to carry novel motif architectures (AATAC and AACAT) that currently lack clinical classification, highlighting the necessity of robust motif decomposition for discovering and cataloguing undocumented sequence diversity (Fig 7c).

### Pathogenic and Biallelic Profiling of RFC1

The *RFC1* locus represents one of the most polymorphic TR sites in the human genome, exhibiting substantial inter-individual variability in both repeat length and motif sequence composition (Weisburd et al., 2026). Biallelic expansions of specific motifs (AAGGG, AAAGG, and ACAGG) are linked to an autosomal recessive syndrome called the cerebellar ataxia, neuropathy, and vestibular areflexia syndrome (CANVAS) (Rafehi et al., 2019). Pathogenicity at the *RFC1* locus is closely tied to the progressive substitution of adenine residues with guanine within the repeat unit; motifs with higher guanine content display an increased propensity to drive disease phenotypes (Wang et al., 2024).

We evaluated *RFC1* variations across a cohort of 498 individuals. Utilizing the LPM tag, we initially identified 30 distinct alleles, varying in both length and sequence composition from 29 unique samples. These alleles were subsequently filtered using motif-specific pathogenic size thresholds: >200 units for the AAGGG motif, and >500 units for the AAAGG configuration (Dominik et al., 2023). Based on these cutoffs, 16 alleles could be classified as truly pathogenic (Table S10). Notably, in sample HG01258, we identified a bi-allelic expansion where the shorter allele carried 423 copies of the AAAGG motif and the longer allele had 580 copies. Applying the motif-specific threshold, only the 580-copy allele was classified as pathogenic. In total, 17 carriers of true pathogenic *RFC1* motif expansions were identified among the 498 individuals (Fig 7d), representing a carrier frequency of 3.4% (Davies et al., 2022).

### Structural and Motif Constraints at FGF14

Among the TR loci examined, *FGF14* emerged as a particularly compelling example of how sequence-level analysis can reveal an interplay between repeat composition and disease susceptibility. An intronic GAA repeat expansion within *FGF14* is causally associated with Spinocerebellar Ataxia Type 27b (SCA27b) (Kytövuori et al., 2025). However, accumulating evidence suggests that interruptions within the repeat tract can exert a protective effect, mitigating pathogenicity even in the presence of substantial repeat expansion. Prior studies have reported that individuals harboring expanded alleles interspersed with alternative motifs (e.g. GGA, GAAGGA), or adenine to cytosine substitutions remain clinically unaffected (Laß et al., 2025; Mohren et al., 2024).

Using the motif decomposition from the DS tag, we observed high motif heterogeneity across the *FGF14* alleles, including a subset of expanded alleles carrying the aforementioned protective interruptions (Fig 7e). Beyond motif composition, ATaRVa revealed a crucial structural distinction between alleles in the normal versus expanded length ranges. Alleles of normal repeat length consistently harbored a non-repetitive 17 bp sequence at the 5’ boundary of the repeat tract, a structural feature which was completely absent in expanded alleles (Fig 7f). A two-sided Mann–Whitney U test confirmed that this difference was highly significant with a p value of 1.2 x 10 ^-69^. This observation supports recently published reports implicating this upstream 17 bp sequence as a *cis*-regulatory feature that constrains repeat expansion, the loss of which acts as a permissive event for the hyperexpansions observed in affected individuals (Prasad G et al., 2025).

Similar to the above examples, we re-evaluated all expanded alleles identified in the 1KGP LRSC cohort using the DS and LPM tags. Following our detailed analysis, we concluded that only 4 genes (*TCF4, RFC1, BEAN1,* and *HTT*) carried truly pathogenic alleles, driven by long, uninterrupted alleles of disease-causing motifs (Fig 8). However, it is important to note that the 1KGP DNA samples are derived from immortalized lymphoblastoid cell lines (LCLs). As LCLs are known to accumulate somatic structural variations and TR instability due to prolonged cell passaging, a subset of these expansions may represent cell line artifacts rather than true variations. Regardless of the biological origin, these findings underscore that total allele length alone is an insufficient diagnostic metric; integrating sequence-level resolution via the DS and LPM tags with interactive profiling in VisuaMiTRa proved essential to eliminate false positives and definitively classify true pathogenic expansions.

**Figure 8:**
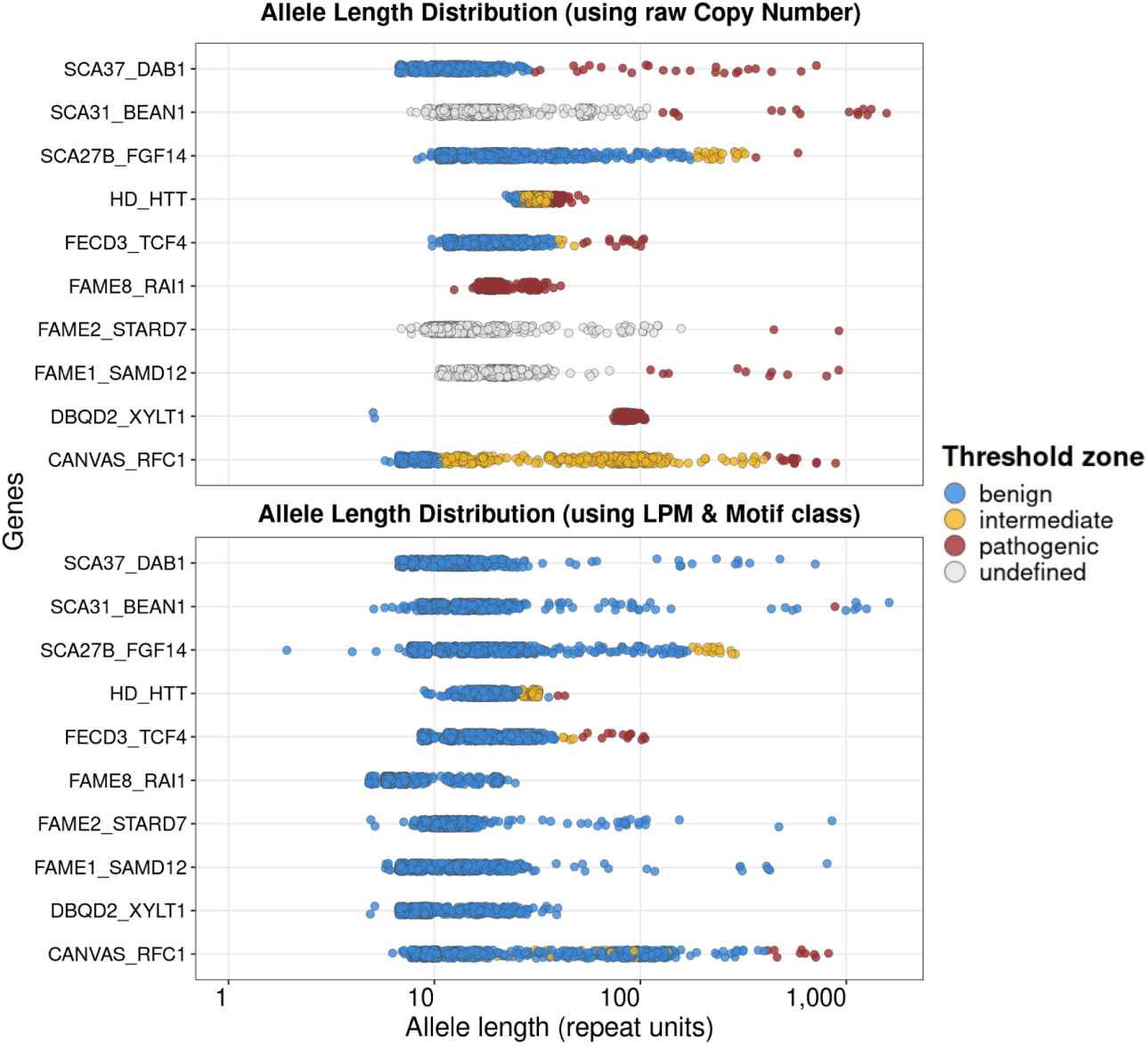
Systematic reclassification of structural false positives using sequence-aware length metrics. Top: Allele length distributions across ten disease-associated loci in the 1KGP LRSC cohort, calculated using raw copy number. Relying exclusively on total allele length classifies numerous alleles across multiple genes as pathogenic (red). Bottom: Re-evaluation of the identical loci applying the Longest Pure Motif (LPM) and target motif class metrics. By quantifying only the maximum uninterrupted run of the disease-associated motif, the vast majority of apparent pathogenic expansions are revealed to be interrupted or structurally benign tracts and are reclassified accordingly (blue).

## Discussion

While the advent of long-read sequencing (LRS) has enabled the study of tandem repeats that were previously intractable, the computational frameworks required to translate this advancement into population-scale biological insights have lagged significantly. Current genotyping algorithms force researchers into a compromise between processing speed, platform flexibility, and genotyping accuracy. ATaRVa resolves this bottleneck by providing a highly scalable, sequencing technology-agnostic framework that combines computational efficiency with highly accurate, structurally aware TR analysis (Table 1).

**Table 1:**
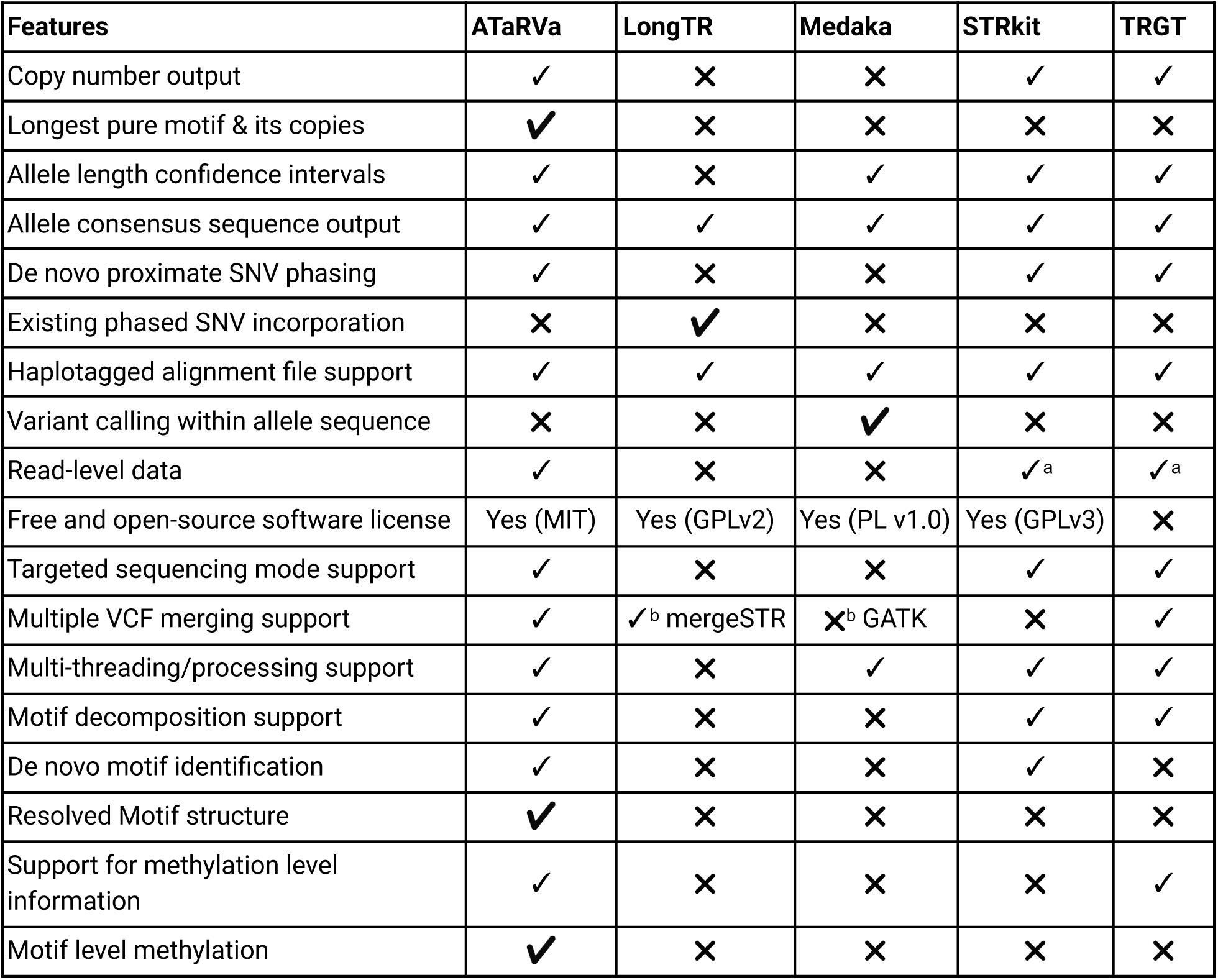
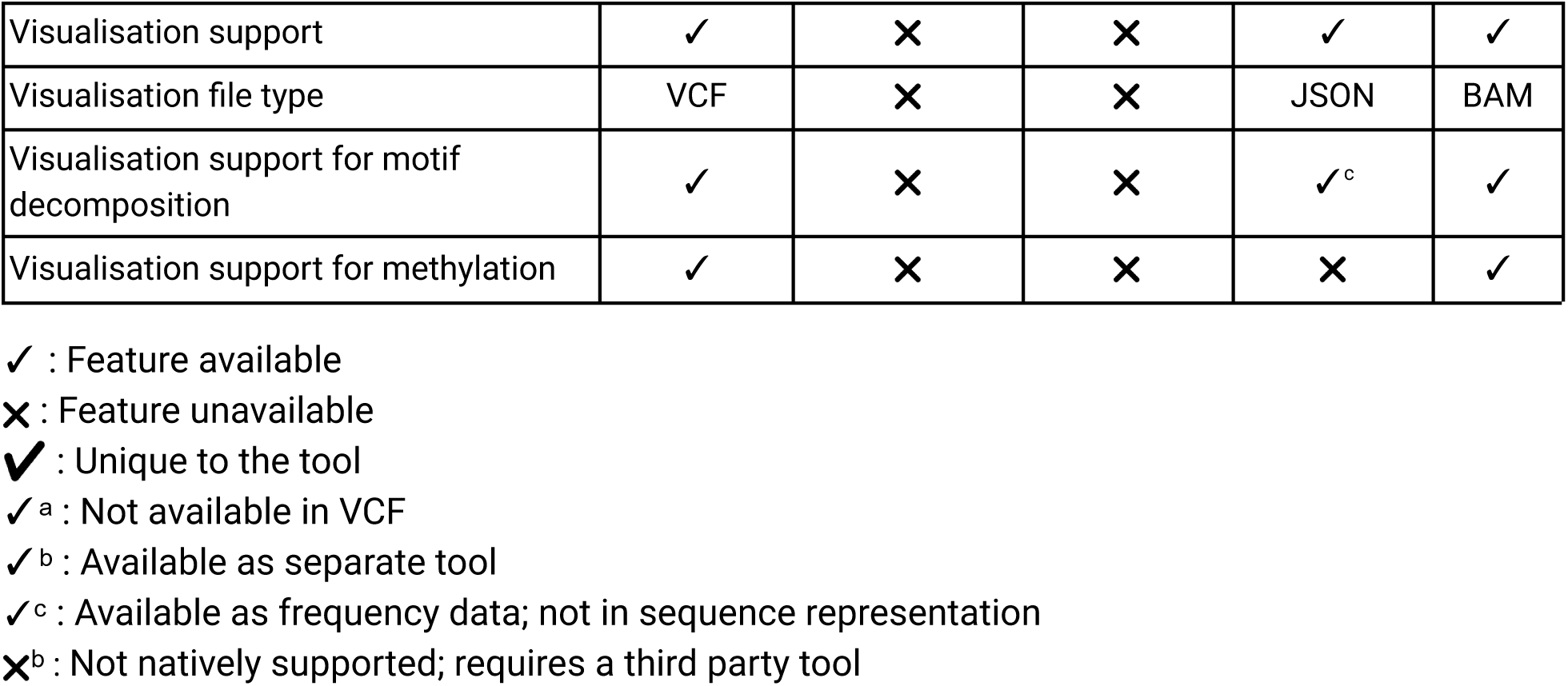
Feature comparison of long-read tandem repeat genotyping frameworks.

A critical limitation in current TR genotyping methods, highlighted by our comparative analyses, is the presence of hidden algorithmic biases. While Medaka Tandem reports high global concordance on ONT data, our zygosity-stratified analysis reveals this is an artifact of a severe bias toward homozygous calls, resulting in a 62.7% error rate for heterozygous loci. Conversely, LongTR exhibits the opposite bias, demonstrating a 27.1% error rate on homozygous loci. In the context of analyzing new genomes or clinical diagnostics, such algorithmic skewing renders these tools unreliable. In contrast, by employing precise flank-realignment and unbiased clustering, ATaRVa delivers highly balanced accuracy across both PacBio and ONT platforms, maintains consistent performance regardless of locus zygosity, and especially outperforms other tools at low sequencing depths.

This balanced performance translates directly into high clinical sensitivity. Across 33 diverse pathogenic expansion samples, ATaRVa provided clinically accurate diagnostic categorizations in 32 cases. While TRGT achieved similar sensitivity owing to its methodological advantage on the PacBio-curated reference datasets, ATaRVa outperformed all other evaluated tools, which consistently underestimated or completely failed to resolve massive expansions. Furthermore, our subsequent application of ATaRVa to the 1000 Genomes long-read cohort demonstrates that clinical TR diagnostics must evolve beyond simple length estimation. As observed in our population-scale analysis, pure length-based screening is highly susceptible to generating structural false positives when biologically benign, interrupted repeat tracts mimic the size of pathogenic expansions. To systematically resolve this, we introduce the Longest Pure Motif (LPM) as a critical new standard metric for TR analysis. By actively quantifying the maximum uninterrupted run of a target motif alongside the complete structural architecture provided by the DS tag, ATaRVa enables researchers to definitively differentiate true pathogenic expansions from complex but harmless alleles. Finally, to facilitate rapid clinical interpretation, the VisuaMiTRa interface enables users to interactively explore these high-resolution motif compositions and their associated epigenetic profiles.

Despite its analytical advantages, ATaRVa operates within several technical boundaries. First, in its default WGS mode, the algorithm primarily relies on aligned read segments; consequently, massive repeat expansions extending into unaligned soft-clipped regions may not be fully recovered, though this limitation is actively mitigated in the `--amplicon` mode. Second, as a unified, platform-agnostic framework lacking deep-learning error correction, resolution of homopolymers is intrinsically tied to raw sequencing quality, resulting in higher performance on PacBio HiFi reads compared to Oxford Nanopore data. Finally, the advanced profiling modules are currently restricted to tandem repeats with motif lengths of up to 10 bp and 5-methylcytosine (5mC) modifications in the CpG context. Expanding the decomposition logic to accommodate larger, more complex VNTR architectures and integrating a wider array of non-CpG methylation marks represent the key areas for future development.

In summary, ATaRVa is a fast, accurate, and structurally aware TR genotyping framework that enables nuanced biological analyses. It delivers the computational throughput necessary to uncover novel TR variations at population-scale, alongside the sequence level resolution required for individual clinical diagnostics. Together, ATaRVa provides the foundation for researchers to systematically investigate the repetitive dark matter of the genome.

## Methods

### Assembly concordance

Genotyping concordance was evaluated using a catalog of 5.5 million isolated tandem repeat (TR) loci derived from TRExplorer v0.2. The HG002 ONT Duplex dataset was obtained from the HPRC, while the ONT Simplex dataset was downloaded from the ONT Open Data repository hosted on AWS. PacBio HiFi datasets for HG002 and five additional samples (HG00438, HG00621, HG00673, HG02622, and HG02630) were obtained from the HPRC (*Human Pangenome Reference Consortium*, n.d.; Liao et al., 2023). The aligned sequencing datasets were genotyped using five tandem repeat genotypers against the TR catalog, producing a VCF file for each sample and tool.

For HG002, the corresponding diploid assembly (HG002 v1.1) was downloaded from the Q100 T2T Consortium, while diploid assemblies for the remaining samples were obtained from the HPRC. The haploid (maternal and paternal) assemblies for each sample were aligned to the GRCh38 reference genome using minimap2 v2.28 with the command: `minimap2 -ax asm5 -t4 GRCh38.fasta.gz sample.fa.gz`. Maternal and paternal allele lengths for each TR locus were subsequently derived from the assembly-to-assembly alignments using a custom script (https://github.com/SowpatiLab/ATaRVa_Manuscript/blob/main/Assembly_allele/aam.py). To ensure high-confidence truth sets, only loci supported by a single contig alignment in both the maternal and paternal haploid assemblies were retained, while all other loci were excluded from downstream analyses.

For each genotyping tool, allele lengths were extracted from the generated VCF files. For ATaRVa, TRGT, and LongTR, allele lengths were obtained directly from the sample genotype fields. For STRkit and Medaka Tandem, allele lengths were calculated from the allele sequences reported in the ALT field. Because LongTR occasionally reports expanded coordinates that include flanking sequence outside the catalog boundaries, coordinate adjustments were applied during evaluation to ensure proper matching with the benchmark catalog.

The set of loci common to both the assembly-derived truth set and the genotyped VCF output of each tool was identified. Genotype concordance was assessed by comparing the reported allele lengths with the corresponding assembly-derived allele lengths. For each locus, the two possible allele pairings between the truth set and the tool-reported genotype were evaluated, and the pairing with the smallest total base-pair deviation was selected. Genotypes were then classified as exact matches, matches within ±1 bp, or mismatches. Overall concordance was calculated as the proportion of loci classified as either an exact match or a match within ±1 bp.

### TR genotyper configuration and execution

To ensure reproducibility and avoid software dependency conflicts, each genotyping tool was executed within its recommended isolated environment. ATaRVa, Medaka Tandem, and STRkit were run within dedicated Python environments, whereas LongTR and TRGT were executed within Conda environments. All evaluated tools were benchmarked using the complete 5.5 million TR locus catalog.

We also attempted to include Straglr and TREAT in the benchmarking analysis. However, both tools exhibited extremely long runtimes without producing results, error messages, or measurable progress when applied to the full catalog. Consequently, they were excluded from further comparisons.

All benchmarking experiments were performed on the same computational cluster running Red Hat Enterprise Linux 8.7 (Ootpa), equipped with an Intel Xeon Gold 6348 processor (2.60 GHz) and 256 GB of RAM.

#### ATaRVa

ATaRVa v0.7.1 was executed using the non-default parameters `--karyotype XY`, `--log`, `--decompose`, and `--methviz`, which were used to treat chromosomes X and Y as haploid in male samples, generate a log file, include motif decomposition information, and record methylation information, respectively. An example command is shown below: `atarva genotype -f reference_genome.fasta -b sample.bam -r TRExplorer.v2.TR.bed.gz -o sample_name.vcf -t 32 --karyotype XY -log --decompose --methviz`

Analyses of pathogenic repeat expansion datasets were performed using the --amplicon mode.

#### LongTR

LongTR v1.2.0 was executed using the non-default parameters `--min-reads 10`, `--min-mapq 5`, `--max-tr-len 25000`, `--min-mean-qual 10`, `--indel-flank-len 10`, and `--haploid-chrs chrX,chrY`. These settings required a minimum of 10 supporting reads for genotyping, a minimum mapping quality of 5, a minimum mean read quality of 10, and allowed genotyping of repeat alleles up to 25 kb in length. In addition, insertions and deletions within 10 bp of the repeat boundaries were considered during genotyping, and chromosomes X and Y were treated as haploid in male samples. The command used was:

LongTR --bam-samps sample_name --bam-libs lib1 --min-mapq 5 --max-tr-len 25000 --min-reads 10 --min-mean-qual 10 –silent --haploid-chrs chrX,chrY --indel-flank-len 10 --bams sample.bam --fasta reference_genome.fasta --tr-vcf sample_tr_calls.vcf.gz --regions TRExplorer.v2.TR.bed

For pathogenic repeat expansion datasets, LongTR was run with --indel-flank-len 20 to maintain consistency with the flank length parameter used in ATaRVa.

#### TRGT

TRGT v5.0.0 was executed using the non-default parameters `--disable-bam-output` and `-k XY`. These settings disabled generation of the output BAM file and specified chromosomes X and Y as haploid for male samples. The command used was:

`trgt genotype -g reference_genome.fasta -r sample.bam -b TRExplorer.v2.TR.bed -o sample_name -t 32 -k XY --disable-bam-output.`

For pathogenic repeat expansion datasets, TRGT was executed using the --preset targeted mode.

#### STRkit

STRkit v0.24.2 was executed using the non-default parameters `--min-reads 10`, `--max-reads 100`, `--flank-size 10`, `--sex-chr`, `--hq`, `--no-tsv`, `--max-mdn-poa-length 10000`, and `--respect-ref`. These settings required a minimum of 10 and a maximum of 100 supporting reads per locus, considered insertions and deletions within 10 bp of the repeat boundaries, treated chromosomes X and Y as haploid in male samples, specified high-quality read data, disabled TSV output generation, skipped partial order alignment (POA)–based consensus generation for loci with a median allele length greater than 10 kb, and constrained tandem repeat coordinates to the reference BED annotations. The command used was:

‘Strkit call sample.bam --min-reads 10 --max-reads 100 --flank-size 10 -p 16 --hq --ref reference_genome.fasta.gz --vcf sample.vcf --loci TRExplorer.v2.TR.bed --sex-chr XY --no-tsv --max-mdn-poa-length 10000 --respect-ref’.

For pathogenic repeat expansion datasets, STRkit was executed using the --targeted mode.

#### Medaka Tandem

Medaka Tandem v2.2.0 was executed using the non-default parameters `--min_depth 10`, `--process_large_regions`, and `--quiet`. These settings required a minimum read depth of 10 for genotyping, enabled processing of tandem repeat loci longer than 10 kb, and suppressed non-essential log messages and warnings. The command used was:

`medaka tandem sample.bam reference_genome.fasta TRExplorer.v2.TR.bed male sample_output --quiet --workers 32 --process_large_regions --min_depth 10`

For the overall concordance analysis using 30× HG002 data and the five additional HPRC samples, a minimum read support threshold of 10 was used for all tools that supported this parameter. TRGT was excluded from this adjustment, as it does not provide an option to modify the minimum read support threshold. For coverage-based analyses on HG002 datasets, the minimum read support threshold was reduced to 5 for all applicable tools to enable evaluation at lower sequencing coverages.

### Assessment of Computational Performance

All performance benchmarking experiments were performed in single-threaded mode to ensure fair comparison across tools. Benchmarking, including runtime and memory consumption analyses, was conducted on a server running Ubuntu 22.04.4 LTS, equipped with an Intel Xeon Gold 6242R processor (3.10 GHz) and 503 GB of RAM. Runtime and memory usage were measured using the UNIX time -v command, and the elapsed (wall-clock) time and maximum resident set size (peak RAM usage) were recorded. For each tool–dataset combination, performance metrics were calculated as the average of five independent runs.

Additional runtime evaluations of ATaRVa features were performed using 34× HG002 datasets containing haplotype-tagged and methylation information, including ONT Simplex data obtained from the ONT Open Data repository and PacBio HiFi data obtained from publicly available PacBio datasets. These analyses were conducted using the default minimum read support threshold of 10 reads.

### Mendelian Concordance Analysis

Whole-genome sequencing (WGS) data for the Ashkenazi trio (HG002, HG003, and HG004), generated using PacBio HiFi sequencing and previously used in the TRGT study, were obtained from NCBI BioProject PRJNA1028149. The datasets were genotyped using ATaRVa, LongTR, STRkit, and TRGT. Medaka Tandem was excluded from this analysis because it is not optimized for PacBio HiFi data.

For each tool, commonly genotyped loci in all three individuals were identified and retained for analysis. At each locus, the offspring alleles were compared against the parental alleles to assess Mendelian inheritance. To determine the most likely inheritance pattern, all possible offspring–parent allele pairings were evaluated, and the combination with the smallest total base-pair deviation was selected. Alleles consistent with inheritance from the parental genotypes were classified as Mendelian, whereas all others were classified as non-Mendelian.

Mendelian alleles were further categorized as exact matches, off-by-1 bp, or off-by-1 motif. Mendelian concordance was calculated as the proportion of Mendelian alleles among all evaluated alleles, with the total number of alleles defined as twice the number of common loci.

The script used for Mendelian concordance analysis is available at https://github.com/SowpatiLab/ATaRVa_Manuscript/blob/main/Mendelian_concordance/mirchi.py

### Motif Decomposition

To characterize the internal structure of such TR locations, ATaRVa provides a motif decomposition feature, enabled by using the `--decompose` flag. This feature enables the user to analyse repetitive stretches of distinct motifs within the allele sequences. ATaRVa generates a condensed representation of the allele sequence indicating distinct repeat motifs and their copy numbers, preserving the intervening sequence interruptions. This feature is developed by leveraging the Ribbit algorithm (Avvaru et al., 2025). The allele sequence is encoded into a bitarray with a 2-bit representation for each nucleotide. The original allele sequence bitarray is aligned with its shifted versions. This shift size ranges from 1 to 10 bps, enabling decomposition up to motif sizes of 10. Individual shift alignments result in bit arrays where matches, indicating shift-size based periodic occurrence of the same nucleotide(s), are represented as 1s and mismatches as 0s. The number and length of matches are compared across all the 10 shifts and optimal decomposition motif size is selected based on the length of consecutive 1s. The motif sequence of the calculated best motif size is identified in the allele sequence using Knuth-Morris-Pratt (KMP) algorithm. Regions containing uninterrupted repeats of the selected motif are condensed into a (MOTIF)n representation. The representative motif is picked after looking at its frequency of occurrence and comparison with its cyclic variations. This motif identification is independent of motif size given in the input repeat catalog. If the majority of the allele sequence cannot be represented by a single motif size, the remaining sequence is iteratively searched for best representative motif size and decomposed with a minimum periodicity of 2 units. Regions are stitched back together along with intervening non-repetitive fragments to reconstruct the full tandem repeat sequence. To provide a clean and meaningful representation of the tandem repeat sequence we have included checks for atomicity of the motif (e.g., converting (AGAG)_2_ to (AG)_4_), checks for consistent repeat class representation, and prioritizing motif size with continuous matches over motif size with more number of fragmented matches.

### Simulated Data for Motif Decomposition

We tested the performance of ATaRVa’s motif decomposition using simulated tandem repeat sequences with diverse sequence complexities (See Data and Code Availability). To generate these datasets, a motif was randomly selected from a predefined list and extended to a random total length ranging from 12 to 200 bp. For the *Pure TR* dataset, these sequences were directly saved into a FASTA file, with corresponding locus and length metadata recorded in a BED file. For the *Interrupted TR* dataset, point mutations were randomly introduced into the extended sequences under two constraints: mutations could not occur within adjacent motif instances, and the overall sequence purity should be maintained at ≥90%. For the *Compound TR* dataset, the pure baseline sequences were interrupted by inserting a second, randomly generated motif extended to a minimum copy number of two. Finally, the *Compound-Interrupted TR* dataset was constructed by introducing these secondary motif interruptions directly into the pre-generated *Interrupted* TR sequences. For each category, a comprehensive BED file was generated to document the sequence length, primary motif type, interruption classifications, and the exact coordinates of all introduced variations.

To access the concordance between simulated data and decomposition script, we implemented a sequence alignment based scoring approach (see Data and Code Availability). For each locus in the simulated data BED (ground truth) and decomposition script BED output were expanded into their full nucleotide representations by repeating every motif according to their copy number, while retaining the boundaries between adjacent motifs as explicit separators. These two expanded sequences were then compared by global pairwise alignment using *Biopython* pairwise2.align.globalms (*Bio.Pairwise2 Module — Biopython 1.76 Documentation*, n.d.). Separator positions were treated as alignment gaps. Alignments were scored with a configurable scheme of match reward, mismatch penalty, and gap-opening and gap extension penalties. For each locus we recorded the raw alignment score and a normalized percent-match, defined as the raw score expressed as a percentage of the maximum attainable score for a perfect alignment of the shorter sequence. Loci were additionally screened for discrepancies in total repeat length and in the number of motif segments, and per-locus scores were aggregated across all loci to summarize overall decomposition accuracy.

Codes used for simulation and testing are available at https://github.com/SowpatiLab/ATaRVa_Manuscript/tree/main/simulation-code.

### Analysis of Samples With Known Pathogenic Expansions

A total of 33 samples carrying pathogenic repeat expansions in *AR, ATN1, ATXN1, ATXN3, C9ORF72, DMPK, FMR1, FXN, HTT, PABPN1,* and *RFC1* genes were collected from multiple public sources (see Data and Code Availability). Twenty-three samples representing all of these disease-associated loci were obtained from PacBio PureTarget sequencing datasets. For these samples, aligned BAM files and corresponding allele length estimates generated using TRGT were available.

Two additional HPRC samples carrying *FMR1* expansions (HG00438 and HG04184) were included. Their allele lengths were obtained from the corresponding TRGT analyses. For HG00438, the aligned BAM file was downloaded directly from the HPRC, whereas for HG04184, PacBio HiFi reads were aligned to the hg38 using minimap2.

Two targeted ONT sequencing datasets generated using adaptive sampling, carrying pathogenic expansions in *DMPK* (NA23265) and *ATXN1* (NA13537) were obtained from NCBI BioProject PRJNA786382. For these samples, pathogenic allele lengths had been determined by molecular diagnostic methods, including repeat-primed PCR (RP-PCR), Southern blotting, or both. Raw sequencing reads were aligned to hg38 using minimap2.

An additional six samples containing expansions in *HTT* (NA13505, NA20253, and NA14044) and *FMR1* (NA13664, NA06896, and NA07537) were obtained from PacBio targeted sequencing datasets, for which hg19-aligned BAM files were available. Pathogenic repeat coordinates for hg38 were obtained from the Adotto catalog (Chiu et al., 2026), whereas hg19 coordinates were provided with the corresponding datasets.

All five genotyping tools were evaluated on the 33 pathogenic samples, except TRGT, which was assessed only on the 31 PacBio datasets. Sensitivity for detecting pathogenic expansions was evaluated by comparing the allele lengths reported by each tool with the corresponding reference lengths provided for each dataset. An expansion was considered successfully detected if the reported allele length was at least as large as the reference allele length. For cases where the reported length was underestimated, the call was still considered concordant if the allele remained within the same clinical classification category (normal, intermediate, premutation, or full mutation/pathogenic), as defined by STRchive for the corresponding disease locus.

### Analysis of 1000 Genomes Project Long-Read Sequencing Consortium Data

The 1000 Genomes Project Long-Read Sequencing Consortium (1KGP-LRSC) is a collaborative effort to resequence a subset of samples from the original 1000 Genomes Project using Oxford Nanopore long-read sequencing technology (Gustafson et al., 2024). The consortium provides aligned sequencing data through a publicly accessible AWS S3 bucket (s3://1000genomes-ont). Leveraging HTSlib’s support for direct streaming from AWS S3, we extracted sliced alignment file with reads aligning at the 66 autosomal pathogenic tandem repeat (TR) loci cataloged in STRchive (Hiatt et al., 2025) (https://strchive.org). Each sample was subsequently genotyped using ATaRVa against the pathogenic TR catalog, and the resulting sample-level VCF files were merged into a single multi-sample VCF using the “ATaRVa merge” command, as follows:

`‘atarva merge -r STRchive-disease-loci.hg38.atarva.bed.gz -i vcf_list.txt -f Homo_sapiens_assembly38.fasta -o 1KGP_merged.vcf -t 12’`

We analyzed allele lengths across 498 samples by comparing the VCF copy number (CN) against pathogenic thresholds from STRchive. True pathogenicity of the expanded alleles was then evaluated using DS and LPM tags, with data extraction and analysis performed via VisuaMiTRA and custom Python scripts.

## Data and Code Availability

ATaRVa is implemented in Python, while VisuaMiTRa is implemented using Python for the backend and React.js for the frontend interface. Both ATaRVa and VisuaMiTRa are freely available through PyPI, and their source code is available under the MIT License on GitHub: https://github.com/SowpatiLab/ATaRVa and https://github.com/SowpatiLab/visuamitra.

Custom code used for analysis in this manuscript is available at https://github.com/SowpatiLab/ATaRVa_Manuscript.

HG002 assembly :

https://s3-us-west-2.amazonaws.com/human-pangenomics/index.html?prefix=T2T/HG002/assemblies/

HG002 PacBio HiFi reads :

https://human-pangenomics.s3.amazonaws.com/submissions/80d00e88-7a92-46d8-88c7-48f1486e11ed--HG002_PACBIO_REVIO/m84039_230117_233243_s1.hifi_reads.default.bam

Ashkenazi trio(HG002, HG003, HG004) PacBio HiFi reads :

https://www.ncbi.nlm.nih.gov/bioproject/PRJNA1028149

HPRC assemblies and PacBio HiFi reads :

https://s3-us-west-2.amazonaws.com/human-pangenomics/index.html?prefix=working/HPRC/

PacBio HiFi samples with repeat expansion:

https://downloads.pacbcloud.com/public/dataset/RepeatExpansionDisorders_NoAmp/

PacBio PureTarget samples with repeat expansions:

https://downloads.pacbcloud.com/public/2024Q4/Vega/PureTargetCoriell24/

Oxford Nanopore targeted sequencing reads with repeat expansion:

https://www.science.org/doi/10.1126/sciadv.abm5386

https://www.ncbi.nlm.nih.gov/bioproject/?term=PRJNA786382

TRExplorer Repeat Catalog:

https://github.com/broadinstitute/trexplorer-catalog/releases/download/v2.0/TRExplorer.repeat_catalog_v2.hg38.1_to_1000bp_motifs.bed.gz

Oxford Nanopore Duplex data for HG002:

https://s3-us-west-2.amazonaws.com/human-pangenomics/index.html?prefix=submissions/0CB931D5-AE0C-4187-8BD8-B3A9C9BFDADE--UCSC_HG002_R1041_Duplex_Dorado/Dorado_v0.1.1/stereo_duplex/

Oxford Nanopore Simplex data for HG002:

https://epi2me.nanoporetech.com/gm24385_ncm23_preview/

ONT simplex and PacBio HiFi haplotagged bams for HG002:

https://downloads.pacbcloud.com/public/revio/2022Q4/WGS-variant-pipeline-analysis/alignments/HG002.GRCh38.haplotagged.bam

https://42basepairs.com/download/s3/ont-open-data/giab_2023.05/analysis/variant_calling/hg002_sup_all/hg002.haplotagged.bam

Python script used to generate Simulated data

https://github.com/SowpatiLab/ATaRVa_Manuscript/tree/main/simulation-code

Simulated data Fasta and BED files

https://github.com/SowpatiLab/ATaRVa_Manuscript/tree/main/manuscript_simulated_data

Python script to assess the concordance in Simulated data

https://github.com/SowpatiLab/ATaRVa_Manuscript/blob/main/simulation-code/alignment-score-cal.py

## Acknowledgements

The authors acknowledge Dr Karthik Bharadwaj Tallapaka and Dr Mohammad Faruq for providing valuable insights in developing the amplicon mode of ATaRVa. We are thankful to the HPC facility of CCMB for providing the computational resources used in this study.

## Author contributions

A.K.S: Software, Formal Analysis, Visualization, Writing - Original Draft; A.S.: Software, Formal Analysis, Visualization, Writing - Original Draft; S.S.: Data Curation, Formal Analysis, Visualization, Writing - Original Draft; S.S.K.: Software, Writing - Original Draft; H.D.: Supervision, Methodology, Writing - Review and Editing; A.K.A.: Conceptualization, Software, Methodology, Writing - Review and Editing; D.T.S.: Conceptualization, Methodology, Supervision, Funding Acquisition, Writing - Review and Editing.

## Funding

This work was financially supported by the grants BT/PR40264/BTIS/137/44/2022 and BT/PR40270/BTIS/137/65/2023 by the Department of Biotechnology, Government of India. H.D. and A.K.A are supported by NHMRC Investigator grant GNT2026126.

## Competing interests

The authors declare no competing interests.

## Notes

https://github.com/SowpatiLab/ATaRVa

### Competing Interest Statement

The authors have declared no competing interest.

### Summary of Updates

This version of the manuscript has been updated with results from ATaRVa v0.7.1 and benchmarked against TRGT, LongTR, STRkit, and Medaka. We showcase new features, including refined motif decomposition and base-level methylation analysis. The results also include analysis of 66 pathogenic tandem repeat locations across 498 samples from the 1KGP ONT consortium. Additionally, we introduce VisuaMiTRa, a new web application designed for visualizing allele-specific motif composition and mean methylation profiles across samples.

