## Supplementary information for "ATaRVa: Analysis of Tandem Repeat Variation from Long-Read Sequencing data"

**for**

### Contents

Supplementary Figures S1 - S12
Supplementary Tables S1 - S10
Detailed Description of ATaRVa and VisuaMiTRa algorithms
References

### Figure S1

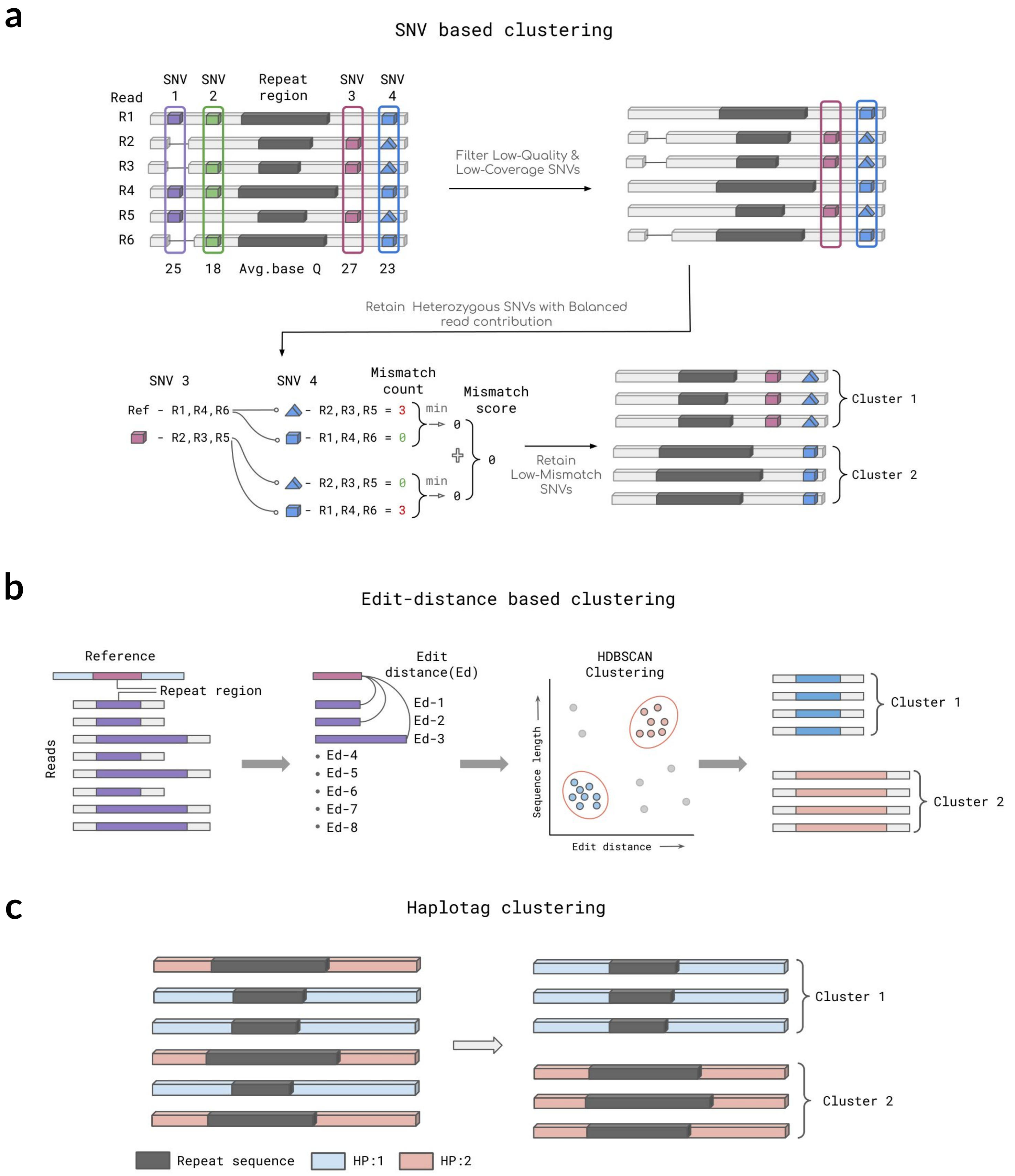


**Figure S1:** Clustering strategies implemented in ATaRVa. a) Overview of SNV-based clustering, illustrating the identification and selection of informative heterozygous SNVs for phasing and clustering supporting reads. b) Workflow of edit distance–based clustering, highlighting the use of repeat sequence edit distance and allele length as input features for HDBSCAN clustering. c) Workflow of haplotag-based clustering, demonstrating the use of pre-phased haplotype information from alignment files to assign reads into haplotype-specific clusters.

### Figure S2

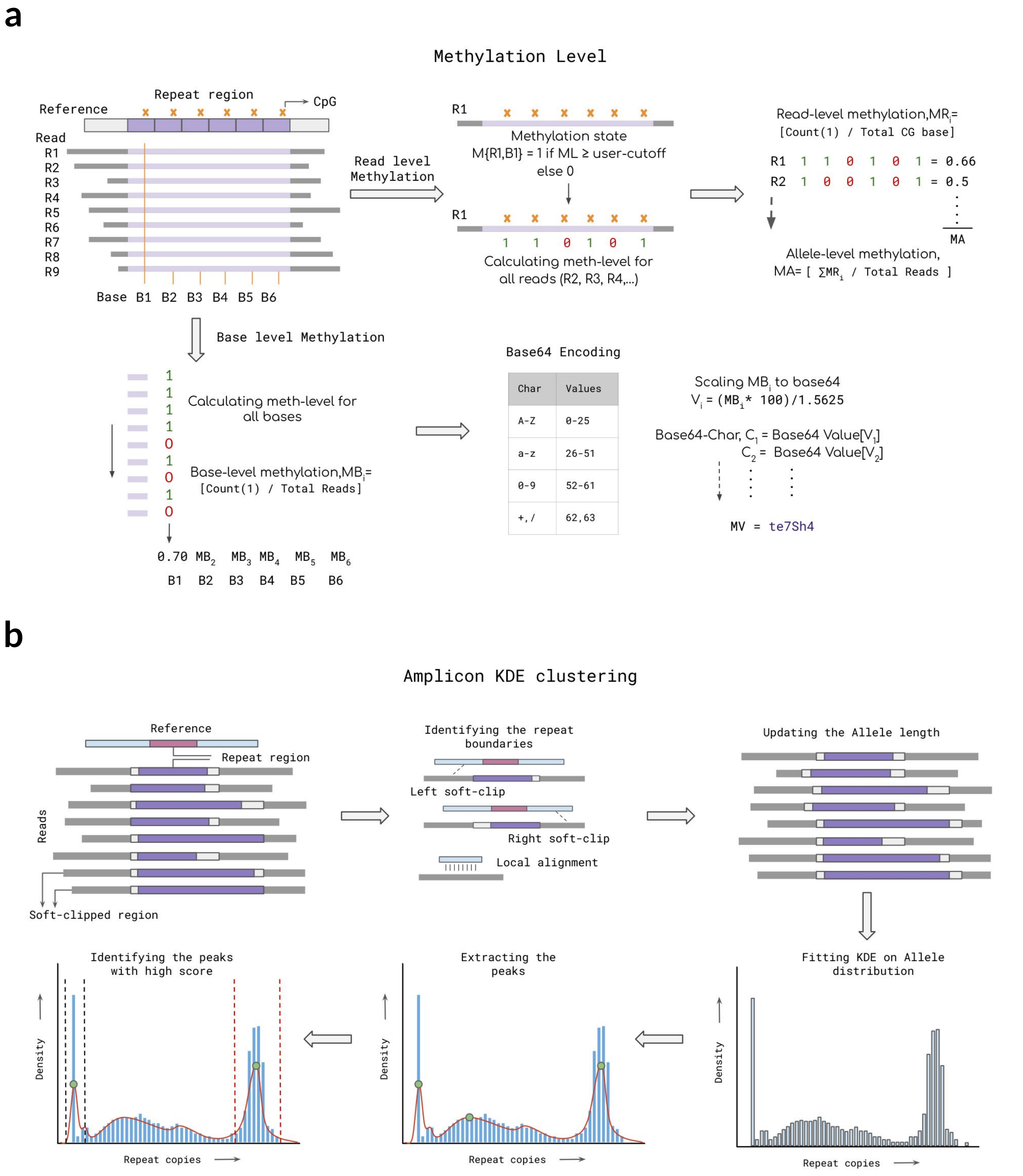


**Figure S2:** a) Overview of CpG 5mC methylation processing, illustrating the calculation of allele-level methylation and generation of per-base methylation profiles for visualization. b) Workflow of KDE-based clustering in targeted sequencing (Amplicon mode), highlighting soft-clipped region processing for repeat boundary extension, allele length estimation, and peak identification for genotype determination.

### Figure S3

a

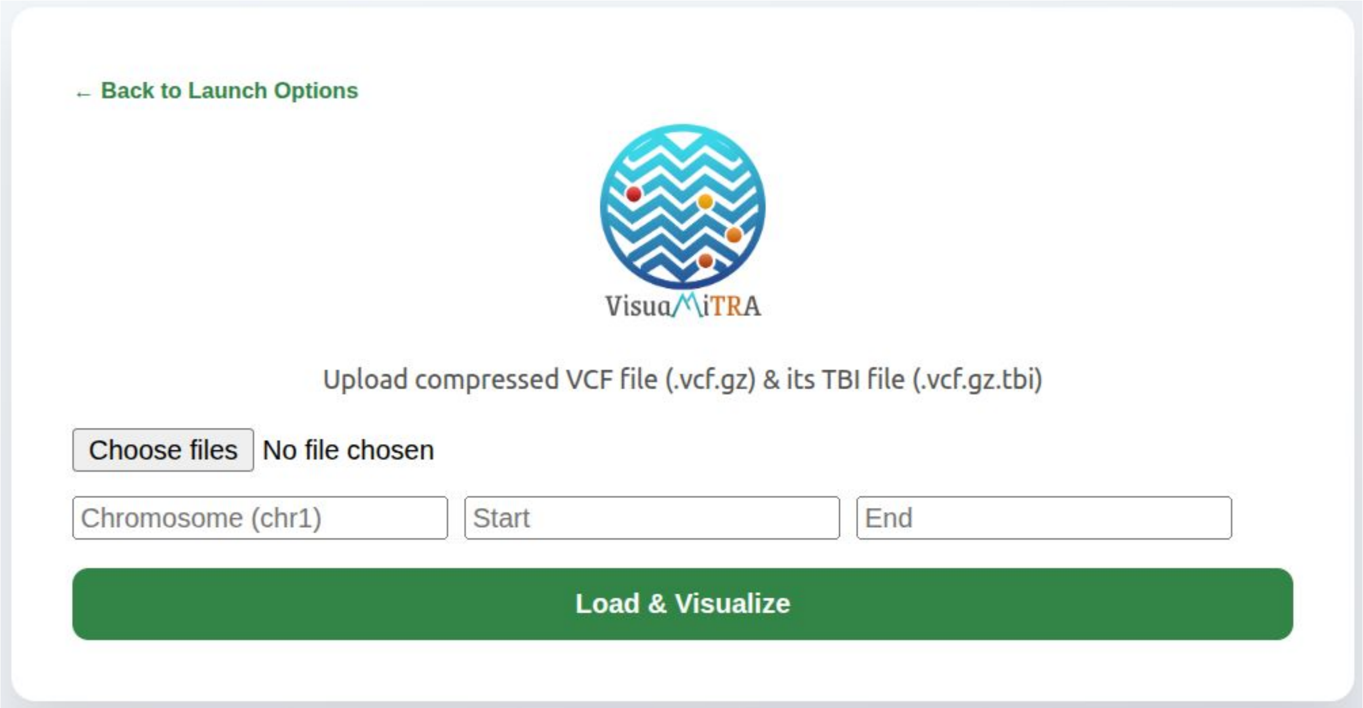


b

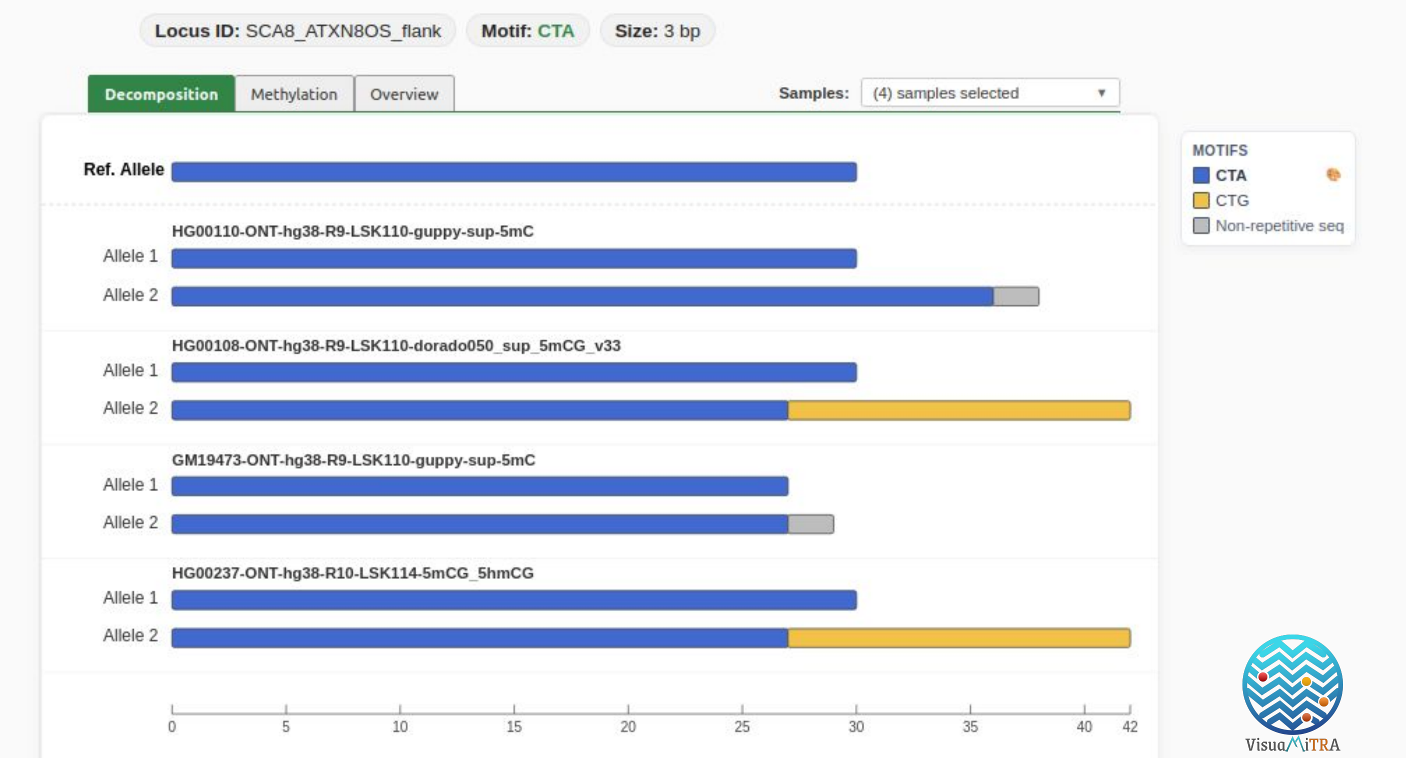


**Figure S3:** VisuaMiTRa User Interface. a) Interface of VisuaMiTRa showing **Browser Mode**, allowing users to upload files through the graphical user interface (single sample or merged VCF) with its index file. b) VisuaMiTRa interface showcasing multisample mode. Comparing motif compositions across 4 samples at locus SCA8_ATXN8OS.

### Figure S4

a

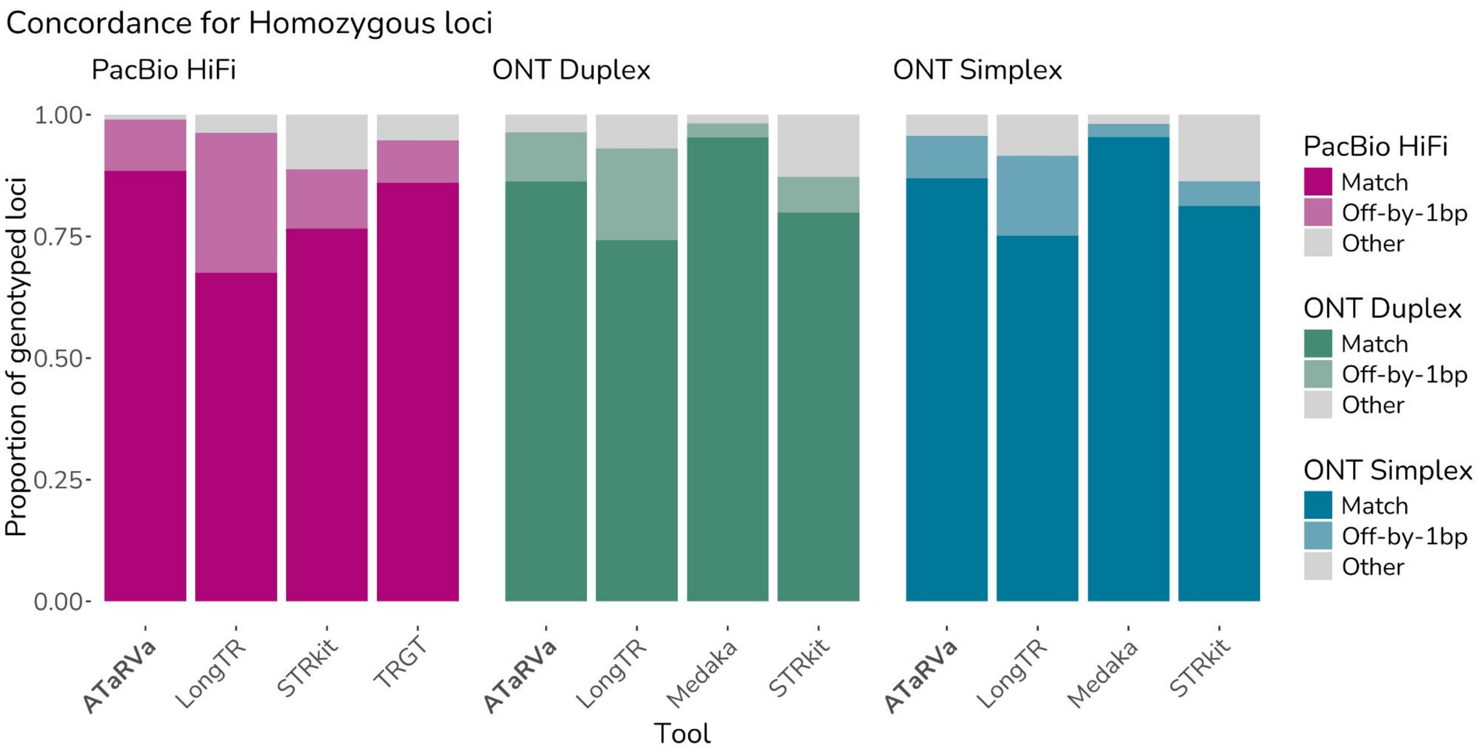


b

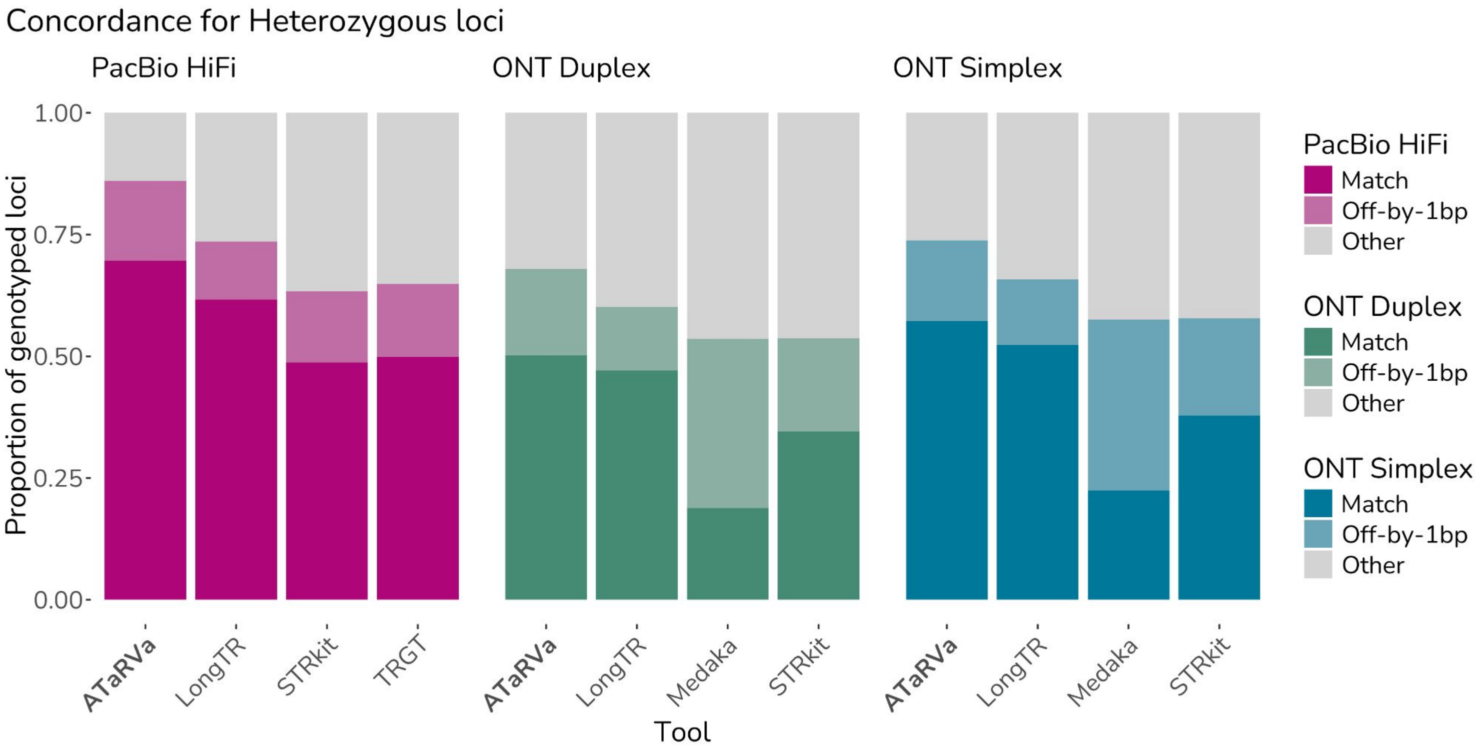


**Figure S4:** Genotyping accuracy and concordance of ATaRVa compared to existing long-read genotypers on the HG002 genome. Overall genotyping performance across three 30× sequencing datasets: PacBio HiFi, ONT Duplex, and ONT Simplex. Stacked bars show the proportion of genotyped loci classified as exact matches to the assembly-derived ground truth (darkest shade), matches within a ±1 bp tolerance (intermediate shade), or mismatches (light grey). Results are stratified by zygosity: a) homozygous loci and b) heterozygous loci.

### Figure S5

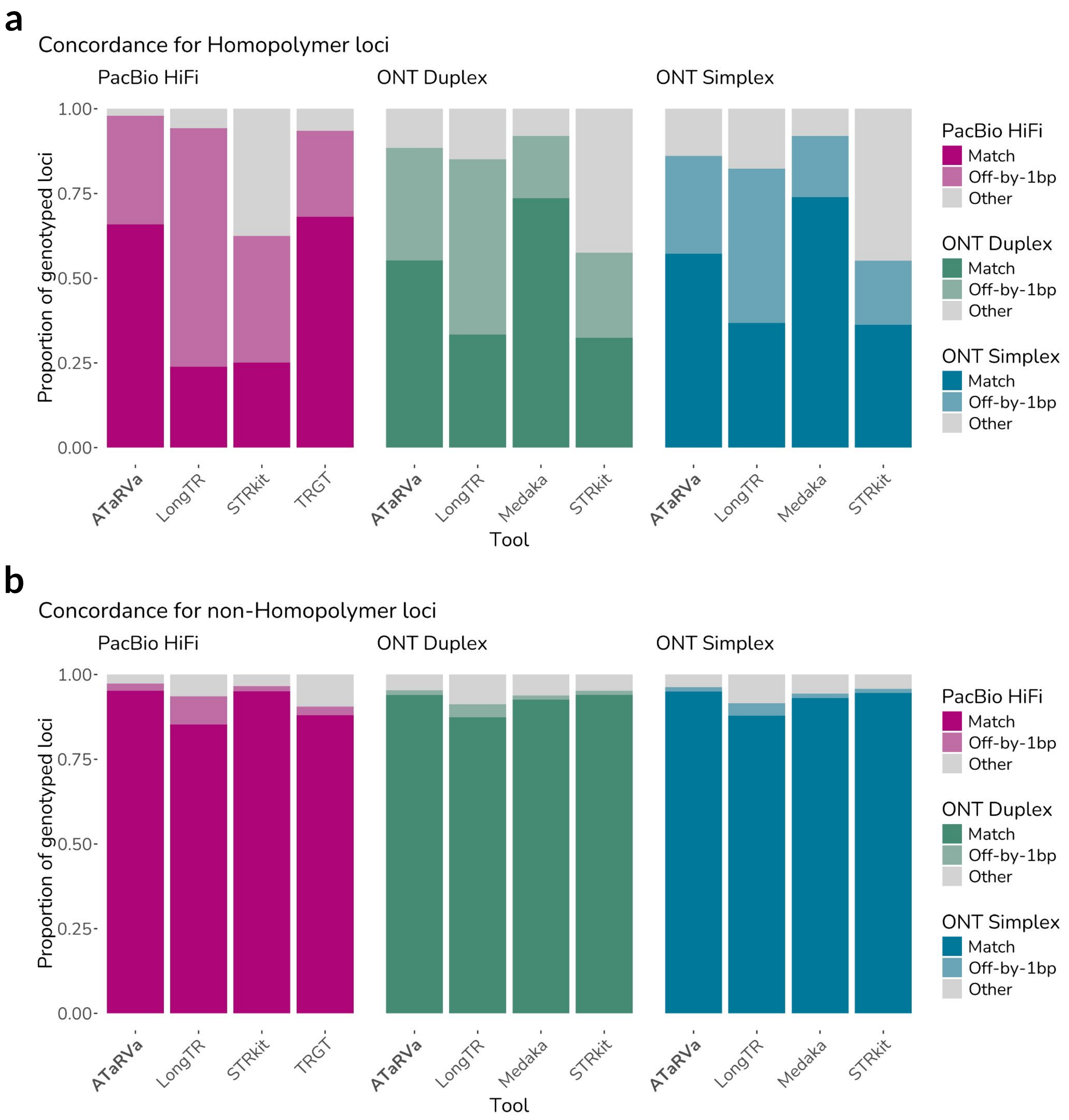


**Figure S5:** Genotyping accuracy and concordance of ATaRVa compared to existing long-read genotypers on the HG002 genome. Overall genotyping performance across three 30× sequencing datasets: PacBio HiFi, ONT Duplex, and ONT Simplex. Stacked bars show the proportion of genotyped loci classified as exact matches to the assembly-derived ground truth (darkest shade), matches within a ±1 bp tolerance (intermediate shade), or mismatches (light grey). Results are stratified by repeat type: a) homopolymer loci and b) non-homopolymer loci.

### Figure S6

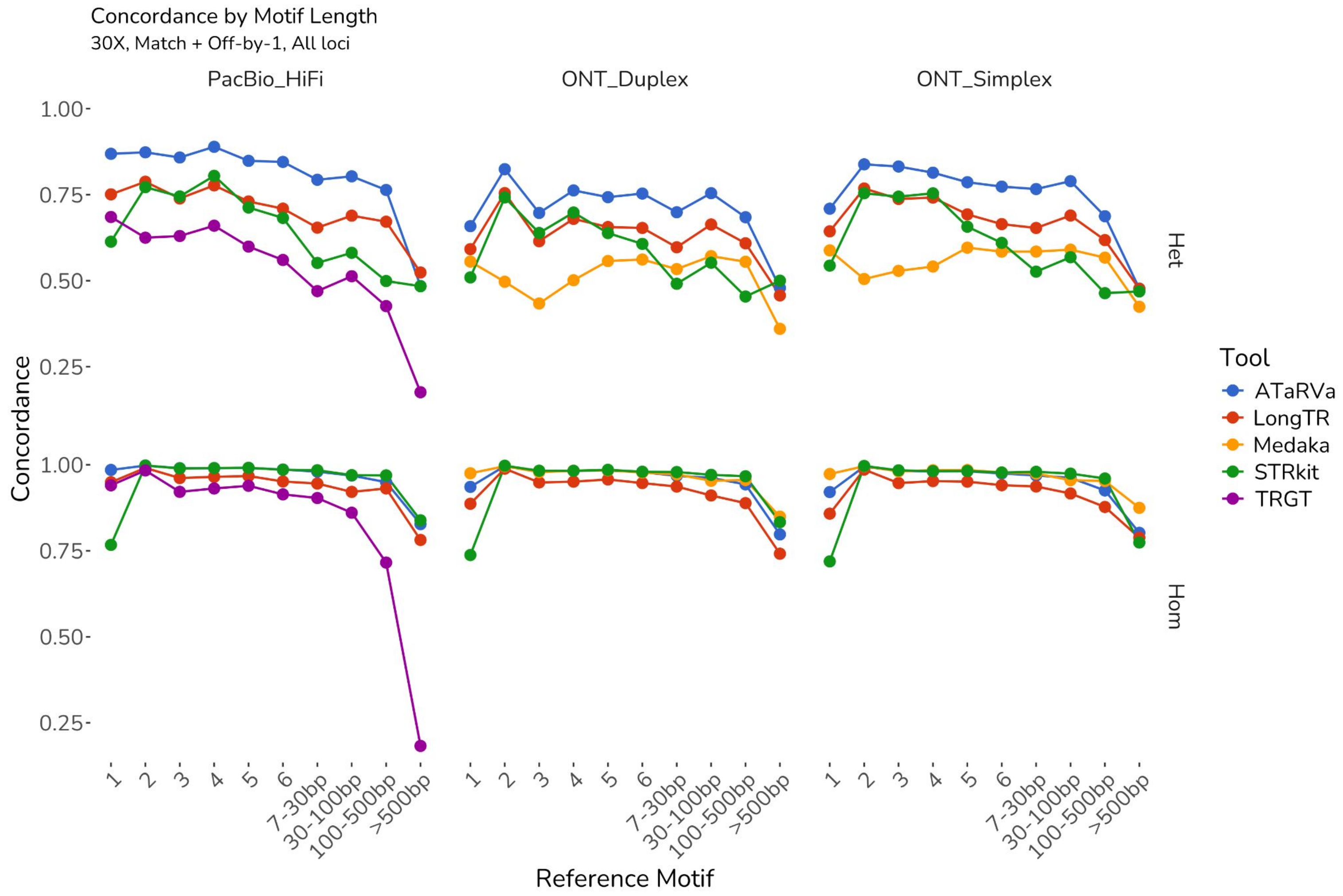


**Figure S6:** Genotyping concordance on the HG002 genome stratified by motif length and zygosity. Overall genotyping concordance, calculated as the proportion of loci classified as either exact matches or matches within a ±1 bp tolerance of the assembly-derived truth set, stratified by reference motif length and locus zygosity. Top row: heterozygous loci; bottom row: homozygous loci.

### Figure S7

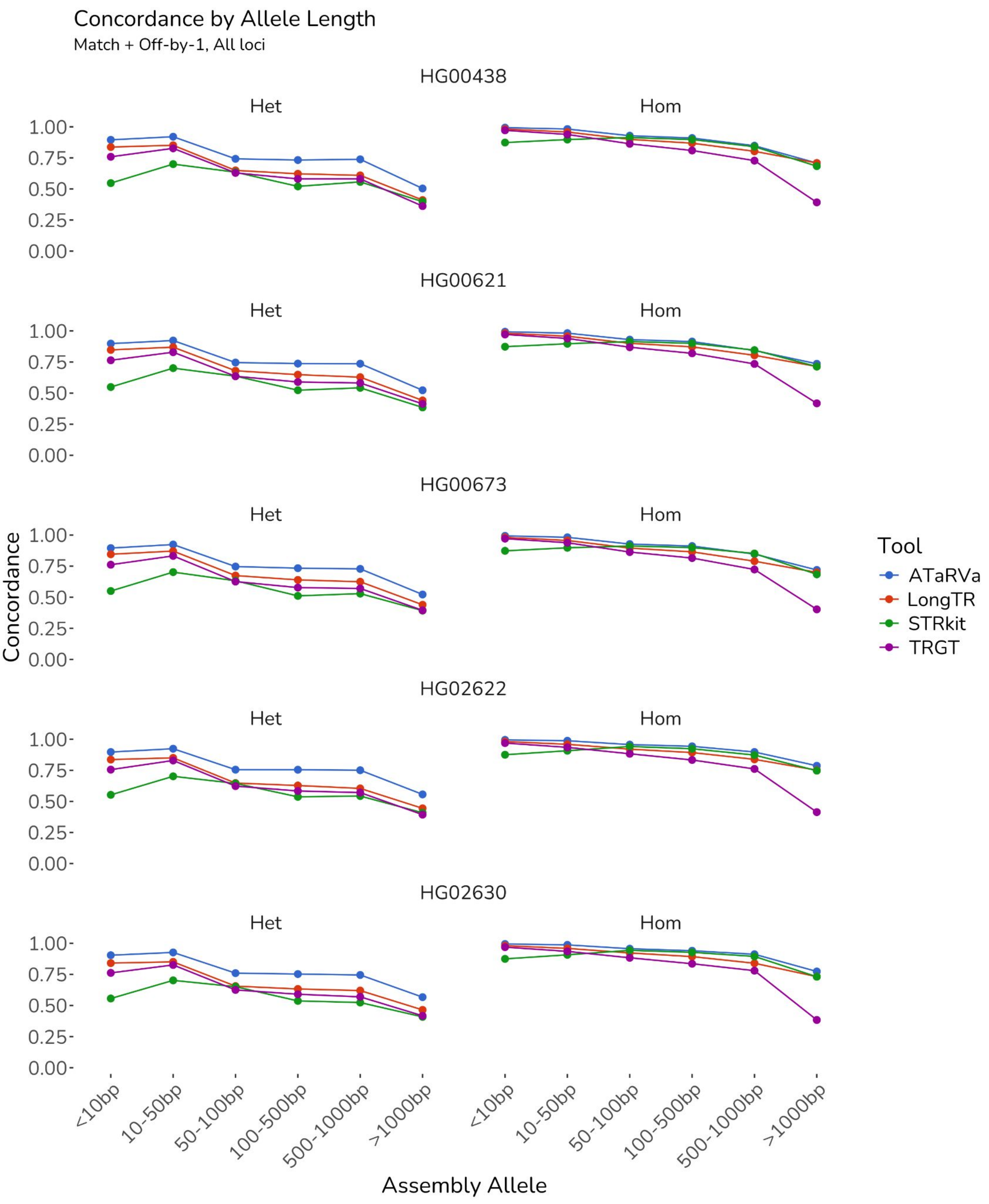


**Figure S7:** Overall genotyping concordance, calculated as the proportion of loci classified as exact match + ±1 bp tolerance of the assembly-derived truth set, across five diverse HPRC genomes sequenced using PacBio HiFi technology. Results are stratified by assembly-derived allele length and locus zygosity.

### Figure S8

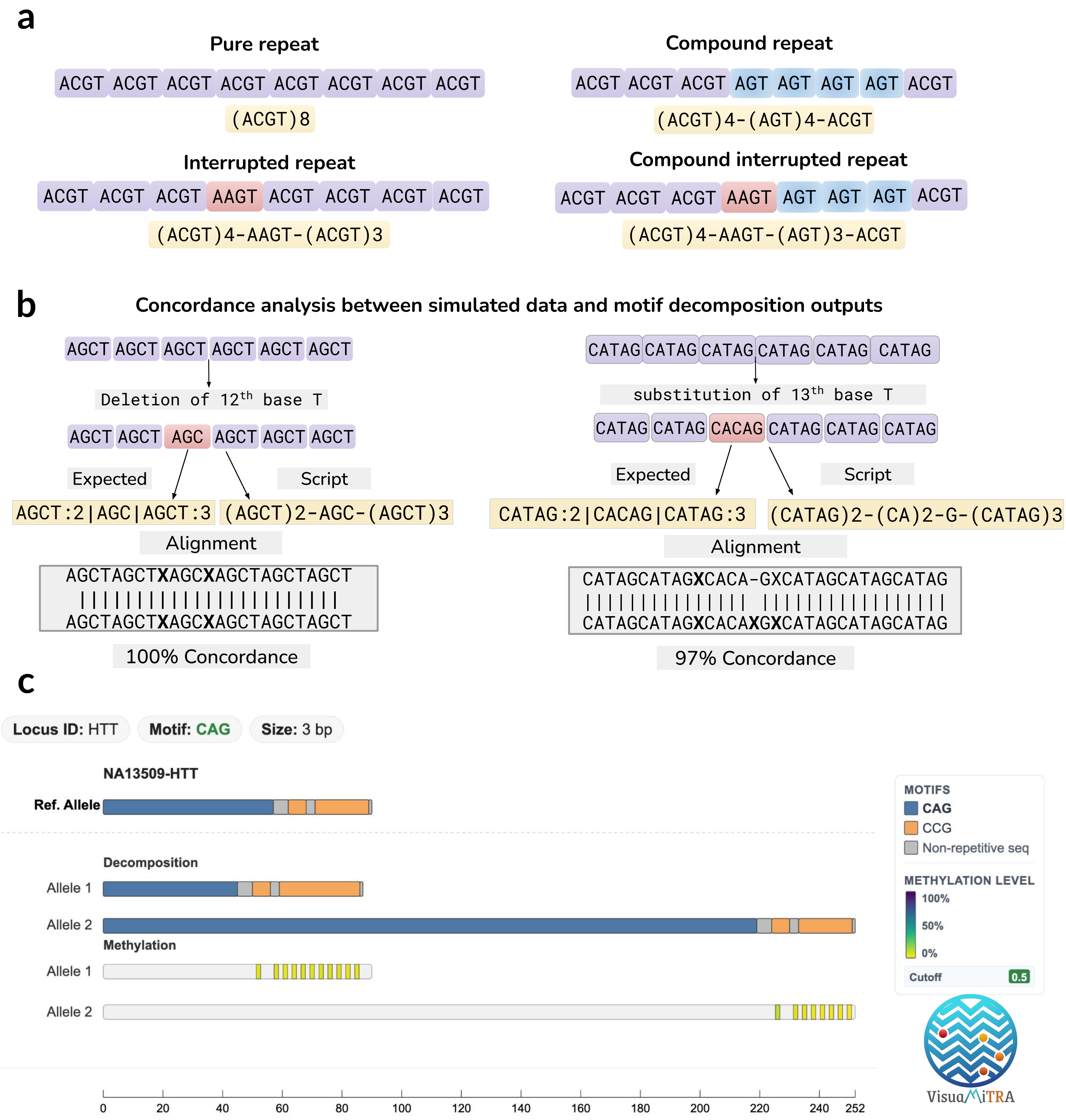


**Figure S8 : (a)** The four distinct categories of simulated data used for validation of the motif decomposition algorithm of ATaRVa. **(b)** Schematic of the alignment-based approach used to determine concordance between the expected motif structure from simulated data and the decomposition script output. **(c)** The VisuaMiTRA interface, demonstrating a representative *HTT* locus example that showcases motif composition, relative allele lengths, and DNA methylation marks.

### Figure S9

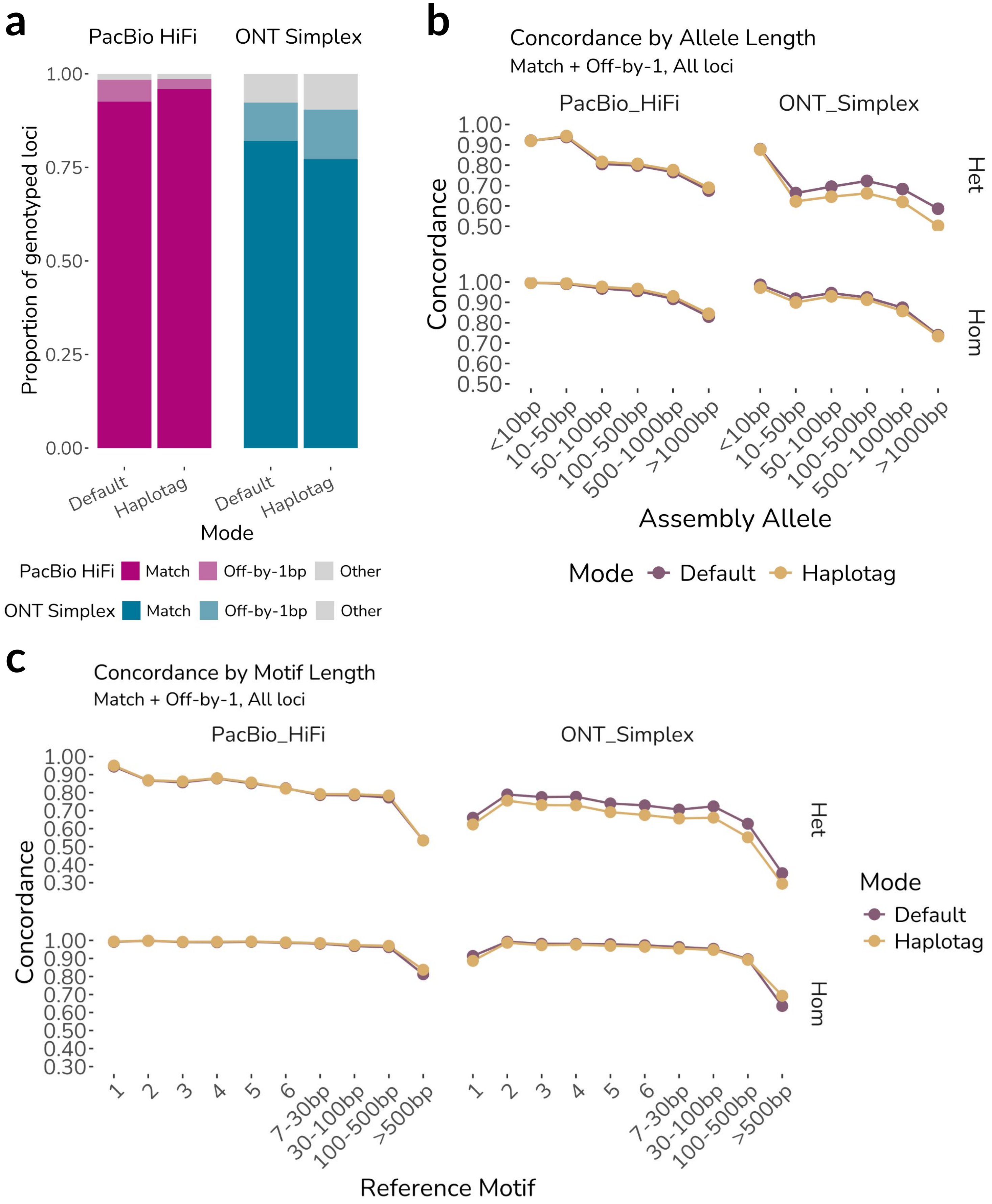


**Figure S9:** Comparison of ATaRVa's default and haplotag modes on the HG002 genome. a) Overall genotyping performance on 30× PacBio HiFi and ONT Simplex datasets. Stacked bars show the proportions of exact matches (darkest shade), matches within ±1 bp (intermediate shade), and mismatches (light grey) relative to the assembly-derived truth set. b) Overall genotyping concordance (exact match + ±1 bp tolerance) stratified by assembly-derived allele length and locus zygosity. c) Overall genotyping concordance (exact match + ±1 bp tolerance) stratified by reference motif length and locus zygosity. Top row: heterozygous loci; bottom row: homozygous loci.

### Figure S10

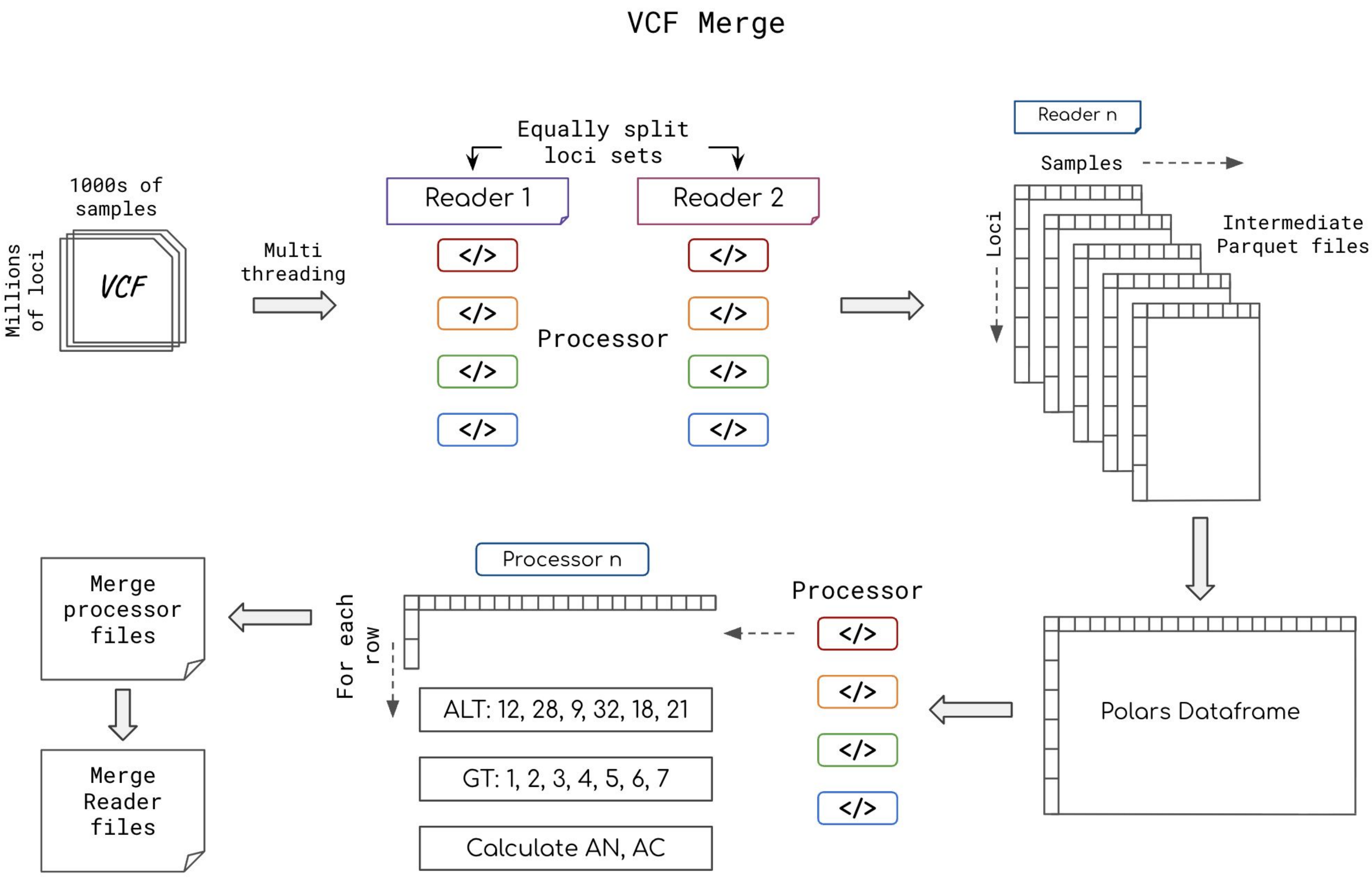


**Figure S10:** Overview of the ATaRVa VCF merging algorithm. Workflow illustrating the scalable merging of large numbers of VCF files containing millions of records, highlighting the multi-threading and multi-processing strategies used to distribute workloads efficiently and generate population-scale merged VCFs.

### Figure S11

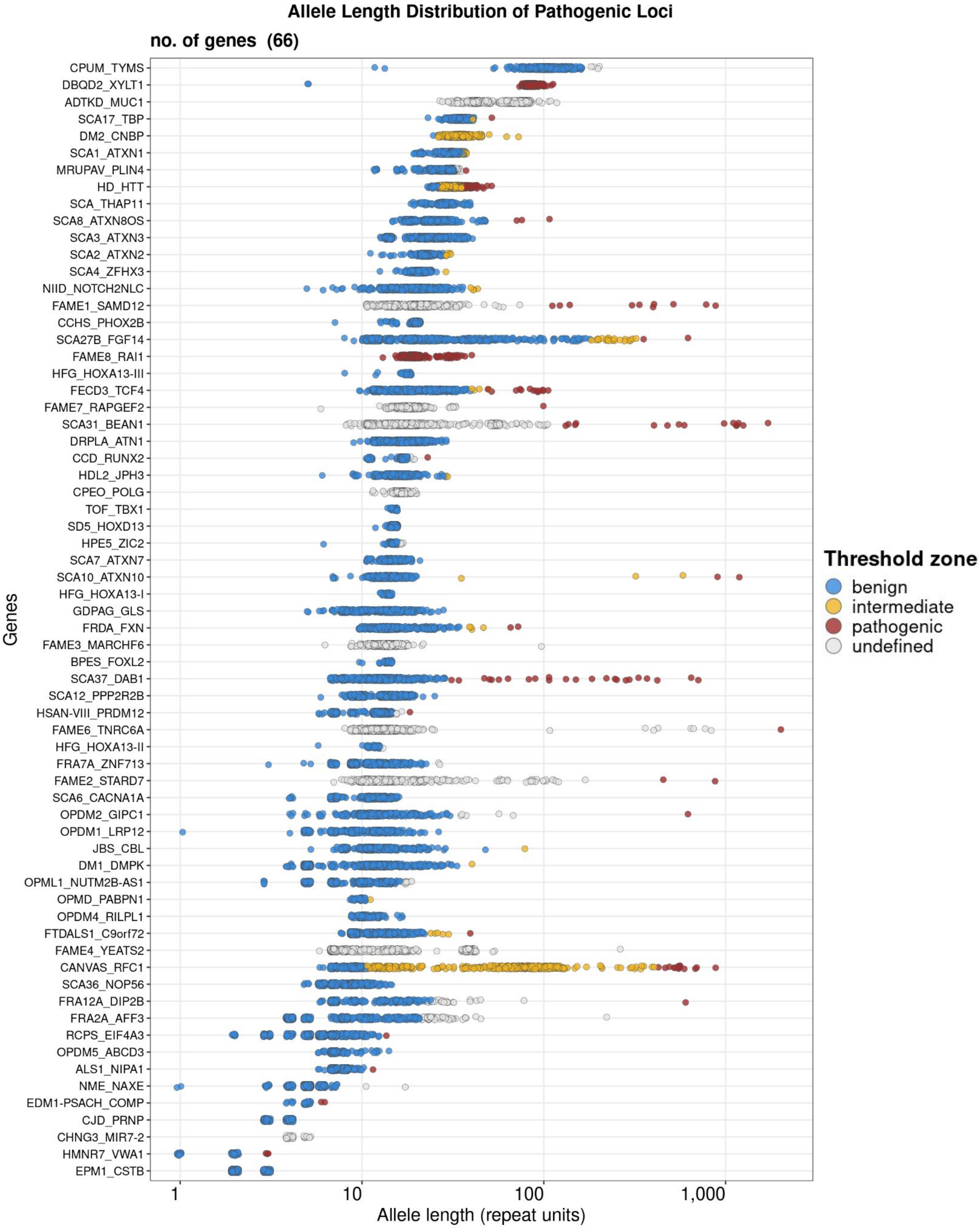


**Figure S11:** Distribution of ATaRVa-derived allele lengths (quantified in repeat units) across 66 autosomal known disease-associated tandem repeats in 498 individuals from the 1KGP Long-Read Sequencing Consortium. Each data point represents an individual allele, color-coded according to published size thresholds: benign (blue), intermediate (yellow), and pathogenic (red). Alleles at loci/sizes lacking defined cutoffs are indicated in grey. Genes are sorted by their median allele length. The x-axis is in log scale.

### Figure S12

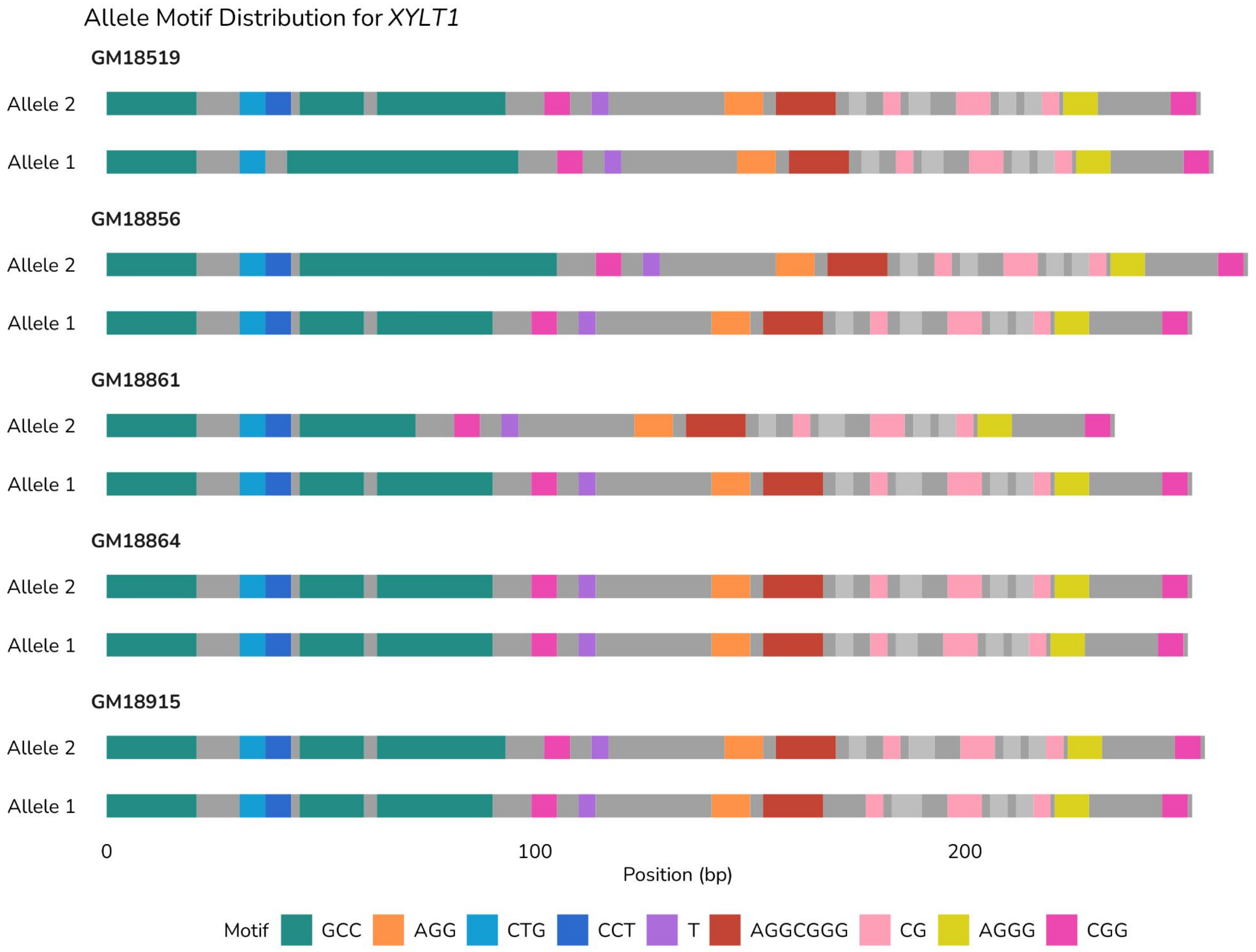


**Figure S12:** Motif composition of *XYLT1* alleles. The green blocks denote GCC repeats; because they do not fully span the alleles, these variants are classified as non-pathogenic.

### Supplementary Tables

**Table S1**: Genotyping rate and assembly concordance of TR genotypers on three datasets of HG002. Bold values indicate the best-performing tool for each metric within the corresponding sequencing platform.

| | Seq. Platform | ATaRVa | LongTR | Medaka | STRkit | TRGT |
|---|---|---|---|---|---|---|
| Number of Genotyped Loci out of 5,599,658 loci | PacBio HiFi | 5497377 (98.17%) | 5501802 (98.25%) | - | 5395954 (96.36%) | **5563509 (99.35%**) |
| | ONT Duplex | 5516108 (98.51%) | 5518144 (98.54%) | **5561834 (99.32%)** | 5409647 (96.61%) | - |
| | ONT Simplex | 5534533 (98.84%) | 5532537 (98.80%) | **5570062 (99.47%)** | 5426467 (96.91%) | - |
| Number of Genotyped Loci out of 5,184,662 loci | PacBio HiFi | 5150887 (99.35%) | 5150766 (99.35%) | - | 4972198 (95.90%) | **5160743 (99.54%)** |
| | ONT Duplex | 5164866 (99.62%) | 5165680 (99.63%) | **5169380 (99.71%)** | 4984451 (96.14%) | - |
| | ONT Simplex | 5168274 (99.68%) | 5168280 (99.68%) | **5170681 (99.73%)** | 4985556 (96.16%) | - |
| Exact Concordance (%) | PacBio HiFi | **86.42** | 66.97 | - | 73.75 | 82.13 |
| | ONT Duplex | 82.32 | 71.34 | **86.96** | 75.21 | - |
| | ONT Simplex | 83.68 | 72.66 | **87.36** | 76.77 | - |
| Concordance with ±1 bp tolerance (%) | PacBio HiFi | **97.53** | 93.94 | - | 86.18 | 91.53 |
| | ONT Duplex | 93.27 | 89.54 | **93.31** | 83.79 | - |
| | ONT Simplex | 93.26 | 88.89% | **93.70** | 83.40 | - |

**Table S2**: Assembly concordance of TR genotypers on three datasets of HG002 in Homozygous loci. Bold values indicate the best-performing tool for each metric within the corresponding sequencing platform.

| | **Seq. Platform** | **ATaRVa** | **LongTR** | **Medaka** | **STRkit** | **TRGT** |
|---|---|---|---|---|---|---|
| Number of Genotyped Loci out of 4,613,712 loci | PacBio HiFi | 4585308 (99.38%) | 4586123 (99.40%) | - | 4460747 (96.68%) | **4594425 (99.58%)** |
| | ONT Duplex | 4596517 (99.63%) | 4597949 (99.66%) | **4600571 (99.72%)** | 4469820 (96.88%) | - |
| | ONT Simplex | 4599544 (99.69%) | 4600279 (99.71%) | **4601749 (99.74%)** | 4470104 (96.89%) | - |
| Exact Concordance (%) | PacBio HiFi | **88.49** | 67.57 | - | 76.60 | 86.07 |
| | ONT Duplex | 86.29 | 74.28 | **95.37** | 79.88 | - |
| | ONT Simplex | 86.95 | 75.12 | **95.39** | 81.25 | - |
| Concordance with ±1 bp tolerance (%) | PacBio HiFi | **98.95** | 96.37 | - | 88.77 | 94.77 |
| | ONT Duplex | 96.40 | 93.11 | **98.20** | 87.22 | - |
| | ONT Simplex | 95.65 | 91.67 | **98.15** | 86.33 | - |

**Table S3**: Assembly concordance of TR genotypers on three datasets of HG002 in Heterozygous loci. Bold values indicate the best-performing tool for each metric within the corresponding sequencing platform.

| | **Seq. Platform** | **ATaRVa** | **LongTR** | **Medaka** | **STRkit** | **TRGT** |
|---|---|---|---|---|---|---|
| Number of Genotyped Loci out of 570,950 loci | PacBio HiFi | 565579 (99.06%) | 564643 (98.90%) | - | 511451 (89.58%) | **566318 (99.19%)** |
| | ONT Duplex | 568349 (99.54%) | 567731 (99.44%) | **568809 (99.63%)** | 514631 (90.14%) | - |
| | ONT Simplex | 568730 (99.61%) | 568001 (99.48%) | **568932 (99.65%)** | 515452 (90.28%) | - |
| Exact Concordance (%) | PacBio HiFi | **69.65** | 62.21 | - | 48.94 | 50.19 |
| | ONT Duplex | **50.23** | 47.55 | 18.93 | 34.64 | - |
| | ONT Simplex | **57.25** | 52.74 | 22.45 | 38.02 | - |
| Concordance with ±1 bp tolerance (%) | PacBio HiFi | **86.04** | 74.23 | - | 63.65 | 65.25 |
| | ONT Duplex | **67.96** | 60.67 | 53.72 | 53.96 | - |
| | ONT Simplex | **73.85** | 66.41 | 57.68 | 58.04 | - |

**Table S4**: Zygosity error rate of TR genotypers on three datasets of HG002. Bold values indicate the best-performing tool for each metric within the corresponding sequencing platform.

| | Tool | PacBio HiFi | ONT Simplex | ONT Duplex | Mean Error |
|---|---|---|---|---|---|
| Homozygous Error | ATaRVa | **11.09%** | 12.22% | 12.63% | 11.98% |
| | LongTR | 31.94% | 24.25% | 25.10% | 27.10% |
| | Medaka | - | **0.78%** | **0.69%** | **0.74%** |
| | STRkit | 19.41% | 10.49% | 12.75% | 14.22% |
| | TRGT | 12.67% | - | - | 12.67% |
| Heterozygous Error | ATaRVa | 7.22% | 9.01% | 10.37% | 8.87% |
| | LongTR | **2.53%** | **2.70%** | **3.18%** | **2.81%** |
| | Medaka | **-** | 62.38% | 63.16% | 62.77% |
| | STRkit | 13.15% | 24.16% | 22.09% | 19.8% |
| | TRGT | 7.63% | - | - | 7.63% |

**Table S5**: Assembly concordance of TR genotypers on three datasets of HG002 in Homopolymer loci. Bold values indicate the best-performing tool for each metric within the corresponding sequencing platform.

| | **Seq. Platform** | **ATaRVa** | **LongTR** | **Medaka** | **STRkit** | **TRGT** |
|---|---|---|---|---|---|---|
| Number of Genotyped Loci out of 1,551,534 loci | PacBio HiFi | 1545773 (99.63%) | 1545595 (99.62%) | - | 1520389 (97.99%) | **1548778 (99.82%)** |
| | ONT Duplex | 1547946 (99.77%) | 1547960 (99.77%) | **1548937 (99.83%)** | 1525310 (98.31%) | - |
| | ONT Simplex | 1548884 (99.83%) | 1548738 (99.82%) | **1549452 (99.87%)** | 1525892 (98.35%) | - |
| Exact Concordance (%) | PacBio HiFi | 65.86 | 23.90 | - | 25.15 | **68.12** |
| | ONT Duplex | 55.25 | 33.38 | **73.69** | 32.44 | - |
| | ONT Simplex | 57.30 | 36.78 | **73.91** | 36.31 | - |
| Concordance with ±1 bp tolerance (%) | PacBio HiFi | **97.97** | 94.30 | - | 62.46 | 93.50 |
| | ONT Duplex | 88.45 | 85.19 | **92.02** | 57.56 | - |
| | ONT Simplex | 86.11 | 82.36 | **92.02** | 55.15 | - |

**Table S6**: Assembly concordance of TR genotypers on three datasets of HG002 in non-Homopolymer loci. Bold values indicate the best-performing tool for each metric within the corresponding sequencing platform.

| | **Seq. Platform** | **ATaRVa** | **LongTR** | **Medaka** | **STRkit** | **TRGT** |
|---|---|---|---|---|---|---|
| Number of Genotyped Loci out of 3,633,128 loci | PacBio HiFi | 3605114 (99.23%) | 3605171 (99.23%) | - | 3451809 (95.01%) | **3611965 (99.42%)** |
| | ONT Duplex | 3616920 (99.55%) | 3617720 (99.58%) | **3620443 (99.65%)** | 3459141 (95.21%) | - |
| | ONT Simplex | 3619390 (99.62%) | 3619542 (99.63%) | **3621229 (99.67%)** | 3459664 (95.23%) | - |
| Exact Concordance (%) | PacBio HiFi | **95.24** | 85.45 | - | 95.16 | 88.14 |
| | ONT Duplex | 93.91 | 87.59 | 92.64 | **94.07** | - |
| | ONT Simplex | **94.98** | 88.02 | 93.12 | 94.63 | - |
| Concordance with ±1 bp tolerance (%) | PacBio HiFi | **97.34** | 93.79 | - | 96.63 | 90.69 |
| | ONT Duplex | 95.34 | 91.41 | 93.86 | **95.35** | - |
| | ONT Simplex | **96.31** | 91.69 | 94.42 | 95.86 | - |

**Table S7**: Mendelian concordance of ATaRVa, LongTR, STRkit and TRGT on the PacBio HiFi data of the Ashkenazi Trio (HG002, HG003, HG004). Bold values indicate the best-performing tool within each category.

| Loci set | Category | | ATaRVa | LongTR | STRkit | TRGT |
|---|---|---|---|---|---|---|
| All loci | Total loci | | 5360022 | 5354912 | 5355237 | 5465087 |
| | Total alleles | | 10720044 | 10709824 | 10710474 | 10930174 |
| | Mendelian alleles | | **10688677 (99.71%)** | 10660651 (99.54%) | 10644687 (99.39%) | 10855114 (99.31%) |
| | Mendelian alleles | Match | **10448867 (97.47%)** | 10380843 (96.63%) | 10296145 (96.13%) | 10636894 (97.32%) |
| | | Off by 1bp | 212341 (1.98%) | 237113 (2.21%) | 260269 (2.43%) | 168523 (1.54%) |
| | | Off by 1motif | 27469 (0.26%) | 42695 (0.40%) | 88273 (0.82%) | 49697 (0.45%) |
| | Non-mendelian alleles | | **31367 (0.29%)** | 49173 (0.46%) | 65787 (0.61%) | 75060 (0.69%) |
| Only non reference loci | Total loci | | 1537877 | 2200253 | 1818634 | 1421645 |
| | Total alleles | | 3075754 | 4400506 | 3637268 | 2843290 |
| | Mendelian alleles | | **3044387 (98.98%)** | 4351333 (98.88%) | 3571481 (98.2%) | 2768230 (97.4) |
| | Mendelian alleles | Match | 2804577 (91.18%) | **4071525 (92.52%)** | 3222939 (88.61%) | 2550010 (89.69%) |
| | | Off by 1bp | 212341 (6.9%) | 237113 (5.39%) | 260269 (7.16%) | 168523 (5.93%) |
| | | Off by 1motif | 27469 (0.89%) | 42695 (0.97%) | 88273 (2.43%) | 49697 (1.75%) |
| | Non-mendelian alleles | | **31367 (1.02%)** | 49173 (1.12%) | 65787 (1.81%) | 75060 (2.63%) |

**Table S8**: Detection of repeat expansions in 33 samples with known pathogenic repeat expansions. *Reported length* represents the expanded allele length (bp) reported by the original study. Values marked with an asterisk (*) are below the reported length but remain within the same premutation or pathogenic range according to STRchive classification criteria.

| Sample | Gene | Reported length | ATaRVa (97%) | LongTR (64%) | Medaka (15%) | STRkit (42%) | TRGT (97%) |
|---|---|---|---|---|---|---|---|
| HM23709 | AR | 147 | 144, 153 | 144, 153 | 69, 69 | 145, 154 | 144, 153 |
| NA13716 | ATN1 | 60, 216 | 60, 216 | 60, 216 | 60, 219 | 61, 220 | 60, 218 |
| ND11494 | C9ORF72 | 48, 4747 | 53, 53 | 18, 48 | - | 18, 18 | 48, 48 |
| NA13536 | ATXN1 | 96, 132 | 96, 132 | 96, 132 | 90, 96 | 97, 133 | 96, 132 |
| NA06153 | ATXN3 | 60, 207 | 60, 207 | 60, 207 | 60, 209 | 61, 211 | 60, 207 |
| NA03697 | DMPK | 36, 1146 | 36, 1146 | 36, 36 | 36, 36 | 37, 36 | 36, 1152 |
| HM03756 | DMPK | 39, 1305 | 39, 1308 | 39, 39 | 39, 39 | 40, 549 | 39, 1317 |
| NA09237 | FMR1 | 2683, 2683 | 2724, 3043 | 2387, 2387* | - | - | 2188, 2713 |
| NA06905 | FMR1 | 69, 237 | 70, 237 | 69, 237 | 69, 240 | 70, 238 | 69, 237 |
| HM06968 | FMR1 | 99, 336 | 100, 337 | 99, 336 | 60, 60 | 102, 341 | 99, 336 |
| NA16237 | FXN | 49, 2095 | 49, 2270 | 49, 48 | 49, 49 | 50, 57 | 49, 2413 |
| NA15850 | FXN | 1802, 2353 | 1956, 3185 | 958, 1007* | - | - | 1848, 3048 |
| NA16202 | FXN | 49, 2388 | 49, 2499 | 50, 51 | 49, 49 | 50, 57 | 49, 2419 |
| NA16212 | FXN | 51, 1523 | 51, 1500 | 53, 54 | 51, 51 | 52, 957 | 51, 1573 |
| NA16215 | FXN | 107, 2963 | 107, 3067 | 110, 111 | 107, 107 | 106, 108 | 107, 2948 |
| ND14442 | C9ORF72 | 12, 4563 | 17, 1296* | 12, 12 | 17, 17 | 18, 18 | 12, 4914 |
| HM23629 | PABPN1 | 18, 27 | 18, 28 | 18, 27 | 18, 18 | 18, 18 | 18, 27 |
| HG01175 | RFC1 | 272, 1948 | 272, 2088 | 272, 272 | 278, 272 | 273, 279 | 2056, 272 |
| HG04228 | RFC1 | 65, 745 | 65, 750 | 65, 65 | 55, 55 | - | 65, 740 |
| NA20752 | RFC1 | 45, 3253 | 45, 3268 | 45, 45 | 45, 45 | - | 45, 3255 |
| NA13509 | HTT | 87, 252 | 87, 252 | 87, 258 | 87, 87 | 88, 255 | 87, 252 |
| NA13515 | HTT | 81, 228 | 81, 228 | 82, 229 | 81, 81 | 82, 232 | 81, 228 |
| HM16212 | FXN | 51, 1426 | 51, 1575 | 52, 53 | 51, 51 | 52, 52 | 51, 1452 |
| NA13505 | HTT | 100, 186 | 99, 186 | 98, 185 | 88, 88 | 102, 102 | 98, 185 |
| NA20253 | HTT | 100, 366 | 99, 363 | 98, 365 | 88, 88 | 102, 102 | 98, 365 |
| NA14044 | HTT | 97, 2155 | 95, 2193 | 95, 2150 | 95, 95 | 99, 1685 | 95, 1879 |
| NA13664 | FMR1 | 92, 159 | 91, 160 | 89, 158 | 92, 158 | 93, 93 | 89, 158 |
| NA06896 | FMR1 | 69, 584 | 70, 416* | 68, 584 | 59, 59 | 69, 596 | 68, 576 |
| NA07537 | FMR1 | 87, 1016 | 86, 1055 | 86, 1031 | 59, 59 | 87, 1022 | 86, 1004 |
| NA13537 | ATXN1 | 90, 180 | 96, 183 | 96, 180 | 90, 96 | 101, 184 | - |
| NA23265 | DMPK | 36, 180 | 36, 221 | 36, 227 | 36, 36 | 36, 228 | - |

| | | | | | | | |
|---|---|---|---|---|---|---|---|
| HG04184 | FMR1 | 90, 174 | 90, 175 | 90, 174 | 60, 93 | 97, 96 | 90, 175 |
| HG00438 | FMR1 | 88, 180 | 87, 180 | 87, 180 | 87, 180 | 88, 181 | 87, 180 |

**Table S9**: Samples with pure expansion of pathogenic motifs

| Gene & Disease | Expanded Alleles Count | True Pathogenic Alleles (based on LPM & motif) | Samples |
|---|---|---|---|
| DBQD2_XYLT1 | 994 | 0 | - |
| FAME8_RAI1 | 124 | 0 | - |
| CANVAS_RFC1 | 22 | 22 | GM20847;HG00253;HG00264;HG01258;HG01342;HG01669;HG01686;HG01816;HG01855;HG01858;HG01862;HG01878;HG02113;HG02220;HG02396;HG03064;HG03698;HG03960;HG03991;HG04026;HG04211 |
| SCA37_DAB1 | 17 | 0 | - |
| SCA31_BEAN1 | 13 | 1 | HG03693 |
| FECD3_TCF4 | 13 | 13 | HG00154;HG00160;HG00233;HG00264;HG00525;HG01669;HG01843;HG02220;HG02351;HG02380;HG03788;HG03910 |
| FAME1_SAMD12 | 9 | 0 | - |
| HD_HTT | 2 | 2 | GM19466;HG02470 |
| FAME2_STARD7 | 2 | 0 | - |
| SCA27B_FGF14 | 2 | 0 | - |

**Table S10**: RFC1 samples carrying alleles expanded beyond the pathogenic threshold

| Sample | LPM-CN | Motif |
|---|---|---|
| HG04189 | 1063 | AAGGG |
| HG01669 | 841 | AAAGG |
| HG04026 | 825 | AAGGG |
| HG02409 | 711 | AAGGG |
| HG01858 | 701 | AAAGG |
| HG02220 | 695 | AAAGG |
| HG01816 | 689 | AAGGG |
| HG01613 | 672 | AAGGG |
| HG01122 | 661 | AAGGG |
| HG04211 | 655 | AAAGG |
| HG00238 | 639 | AAAGG |
| HG01342 | 611 | AAAGG |
| HG03698 | 593 | AAAGG |
| HG01258 | 580 | AAAGG |
| GM20847 | 515 | AAAGG |
| HG01862 | 358 | AAGGG |
| HG01855 | 315 | AAGGG |

### Detailed description of ATaRVa algorithm

#### Parsing data from input files

ATaRVa begins by preprocessing the region BED file to determine the overall start and end positions of the repeat regions in each chromosome. If multiprocessing is enabled, the regions are split into equal chunks and assigned to different processors. Unlike other existing long-read genotyping tools, as default ATaRVa iterates through the alignment file read by read, rather than region by region. During this process, each read is either processed or skipped based on its mapping quality (minimum of 5 by default) and whether it fully encloses at least one repeat region.

The alignment annotations of each read, either the CS tag or the CIGAR string together with the MD tag, are parsed to extract the read-specific start and end coordinates of enclosed repeat loci, single-nucleotide variant (SNV) positions their corresponding alternate alleles and base-calling Phred quality scores (minimum of 20 by default), and deleted reference positions. This information, along with detected base modifications and their associated probabilities, is stored in a read-level data structure using custom read IDs as keys. At this stage, a second primary data structure containing SNV-level information is updated using the variants extracted from each read, with reference genomic positions serving as keys. Following the processing of each read, locus-specific information is incorporated into a global repeat-locus data structure indexed by locus coordinates. This locus-level structure stores supporting read identifiers, allele sequences including flanking regions, and base methylation information within the repeat region.

As reads are processed in genomic order, a repeat locus is considered complete and eligible for genotyping once the start position of the current read exceeds the end coordinate of the locus, indicating that all supporting reads for that locus have been encountered. The accumulated SNV information is then used to cluster reads into haplotypes, and a consensus allele sequence is generated for each haplotype from the corresponding set of supporting reads and the average base methylation of consensus is calculated from all reads.

##### Processing of methylation information:

The methylation information, including the position, strand, and probabilities of modified bases, is extracted from the MM and ML tags of the alignment file for each repeat-spanning read.

The ML tag encodes modification probabilities as integer values ranging from 0 to 255. These values are converted into probabilities and subsequently transformed into binary methylation states: 0 for unmethylated and 1 for methylated, based on a user-defined methylation probability cutoff. By default, a cutoff of 0.5 is used, where probabilities $\geq 0.5$ are considered methylated and probabilities $< 0.5$ are considered unmethylated. Binary methylation values are then assigned to each modified base.

Simultaneously, the mean methylation level for each read is calculated as the average methylation state across all modified bases and stored together with the per-base methylation states. After clustering the reads into two haplotypes, the mean methylation level for each cluster is calculated by averaging the read-level methylation values of all supporting reads within that cluster (Fig S2a).

For visualization of methylation patterns across both alleles, per-base methylation levels are calculated and stored in the VCF as Base64-encoded values. Initially, all CpG positions are extracted from the consensus sequence of each allele and from their corresponding supporting reads. The methylation states of CpG bases from each read are then mapped according to their alignment with the consensus sequence, and the average methylation level for each CpG position is calculated. Finally, these per-base methylation levels, ranging from 0 to 1, are converted into Base64-encoded representations.

The methylation cutoff value influences the assignment of the methylation state for each modified base. All methylation probabilities above the cutoff are considered methylated, while probabilities below the corresponding opposite threshold are considered unmethylated. For example, if the specified cutoff is 0.8, modified bases with probabilities ≥ 0.8 are assigned a value of 1 (methylated), whereas probabilities ≤ 0.2 are assigned a value of 0 (unmethylated). All subsequent steps are then performed as described previously.

#### Genotyping in default mode

In WGS mode, all stored allele lengths in the repeat locus data are updated based on the presence of extended repeat sequences within the flanks, achieved by performing a local realignment between the inserted sequence in the flanking region and the reference repeat region. Once the allele sequence and lengths are updated, the stored methylation information such as base positions and their probabilities is processed to calculate the mean methylation within the updated boundaries of the repeat region for each read and followed by a clustering process to phase these reads into two haplotype sets. There are three types of clustering methods in WGS mode: SNV-based clustering, edit-distance based clustering and haplotag clustering for phased bam.

##### SNV-based read phasing:

Using multiple informative heterozygous SNVs identified across the supporting reads, ATaRVa phases reads into two haplotype groups based on concordant allele-specific read sets (Fig S1a). Prior to this, ATaRVa iterates through each recorded SNV position and examines all reads spanning that site. Reads lacking a recorded SNV at the position are assigned the reference allele, provided that the position is not encompassed by a deletion in the read. This process generates allele-specific read sets for each informative SNV, which are subsequently used for haplotype phasing.

Then, SNVs that are within the user-defined distance on either flank of the repeat locus and with a minimum coverage of five reads or 60% of total coverage are retained. A progressive filtering strategy is applied to identify informative heterozygous SNVs for haplotype assignment. Initially, SNV positions are retained if the two most-supported alleles each contribute 30-70% of the total read support. If no qualifying SNVs are identified, the criteria are relaxed to retain positions with allele having read contributions of 25-75%. If necessary, a final filtering step further relaxes the thresholds to 20-80%, thereby maximizing the likelihood of identifying informative variants while maintaining robustness to sequencing noise and allelic imbalance.

ATaRVa proceeds with haplotype phasing when at least a user-defined minimum number of informative SNVs is identified. When the number of informative SNVs exceeds this threshold, the highest-coverage SNVs are selected for phasing. For each selected SNV, only the two most-supported alleles are retained, and the corresponding allele-specific read sets are constructed. The selected SNVs are then evaluated in descending order of read coverage. Pairwise comparisons are performed between the allele-specific read sets of all SNV pairs, and a mismatch score is calculated to quantify the degree of discordance between their allele assignments. For a given SNV pair, the read set supporting the first allele of SNV1 is compared against both allele-specific read sets of SNV2, and the smaller of the two intersection sizes is retained. The same procedure is repeated for the second allele of SNV1. The sum of these two minimum intersection values is defined as the mismatch score for the SNV pair, with lower scores indicating greater concordance between the two SNVs. This procedure is repeated for all pairwise combinations of selected SNVs. An SNV pair is considered highly informative if it exhibits at least two pairwise comparisons with a mismatch score of zero, indicating complete agreement in allele-specific read assignments. If no such pair is identified, each SNV is assigned a score equal to the sum of its two lowest mismatch scores across all pairwise comparisons. The SNV exhibiting complete concordance with at least two other SNVs (mismatch score = 0) is selected as the seed for haplotype clustering. If no SNV meets this criterion, the SNV with the lowest aggregate mismatch score is chosen instead.

Once an SNV is selected, its pairs are sorted in ascending order based on their initial mismatch scores. Clustering of reads begins by assigning the reads corresponding to the two nucleotides of the selected SNV into two different clusters. Subsequently, the reads of other SNVs, in the sorted order, are assigned to either cluster. For an allele in a successive SNV to be clustered into one of the two clusters, the percentage of common reads between the SNV and the assigned cluster must exceed 70%, and the overlap with the opposite cluster must be below 5%. Finally, the combined number of reads in both clusters must meet the user-defined threshold for the total number of supporting reads in order to be considered as phased alleles. After clustering the reads, the allele for each corresponding read within each cluster is retrieved from the repeat locus data.

If SNV-based phasing cannot be performed due to insufficient informative variants or failure to meet user-defined quality and support thresholds, reads are phased based on the edit distances of allele sequences from the reference.

##### Edit-distance based read phasing:

In the edit-distance-based phasing approach, allele lengths supported by a single read are initially excluded. However, singleton alleles located within 10% of the length of another observed allele are retained and merged back into the primary allele set. A minimum of three supporting reads is required for clustering; loci failing to meet this threshold are excluded from genotyping.

For each retained read, the edit distance between the observed repeat sequence and the reference repeat sequence is calculated using StringZilla (Vardanian, 2020/2026). The resulting edit distances, together with allele lengths, are used as two-dimensional features for HDBSCAN clustering (*HDBSCAN*, n.d.). The minimum cluster size is set to the greater of two reads or 10% of the total supporting reads. Following clustering, up to two allele clusters are retained based on cluster support, corresponding to the clusters with the largest numbers of assigned reads (Fig S1b).

##### Haplotag based read phasing:

This method utilises the haplotype information available in the phased alignment file by storing the haplotype tag for each read (Fig S1c). Once all supporting reads for a given locus are extracted, at least 85% of the total reads must contain haplotype information for the locus to be genotyped using this approach. Since the haplotype tag can have a value of either 1 or 2, reads sharing the same haplotype value are clustered together.

To use this method, the user must enable the " - - haplotag" flag. This mode allows faster processing because it avoids storing SNV information for reads that already contain haplotype tags. Only reads lacking haplotype information are processed for SNV extraction. If the proportion of haplotagged reads does not meet the minimum 85% threshold, the extracted SNV information is used for clustering instead. If SNV-based clustering also fails, the method falls back to edit distance–based clustering.

#### Genotyping in Amplicon mode

In amplicon mode, ATaRVa analyzes unaligned soft-clipped regions of reads that do not fully span the repeat locus and its user-defined flanks (Fig S2b). A default 20 bp (or user-defined) reference flank sequence is extracted from the uncovered flanking side and searched against the soft-clipped region using local alignment. If at least 70% of the reference flank aligns to the soft-clipped sequence, the match is accepted as a valid repeat boundary, and the sequence bounded by the identified flanks is extracted as the repeat region.

Simultaneously, read-level, locus-level, and methylation-related information are extracted and stored, while SNV-related information is omitted. In amplicon mode, all SNV-based phasing is disabled, genotyping is performed solely based on allele length.

After all supporting reads are processed, allele lengths are converted to repeat copy numbers and the resulting distribution is modeled using kernel density estimation (KDE) (*KernelDensity*, n.d.). A Gaussian kernel is employed with a KD-tree implementation, the Minkowski distance metric, and a bandwidth dynamically estimated from the underlying data distribution. A modified Scott's formula is used to estimate the bandwidth:

Bandwidth, $h = 0.5 * \sigma n^{-1/5}$

The fitted KDE is evaluated on 1,000 equally spaced grid points, with the range extending 50 points below the minimum value and 50 points above the maximum value of the data distribution. These additional boundary points are included to ensure proper representation of the Gaussian curve, particularly when peaks occurred near the extremes of the distribution. Using the density values obtained across these 1,000 grid points, peaks in the distribution are identified.

From the multiple detected peaks, a maximum of two peaks are selected based on a custom score calculated for each peak. The scoring formula is defined as follows:

Score, $S = \frac{P^2 . A}{(Wt + \epsilon)^2} \times Qk$

Where,
P = prominence of the peak
A = Area under the peak
Wt = Width of the peak at 80% height from base
$Q_k$ = Skewness-based quality weight for each peak
$\epsilon$ = 1e-8

The skewness-based quality weight ($Q_K$) is a non-linearly scaled score calculated for each peak based on its relative asymmetry using a sigmoid function. Initially, the skewness (K) of each peak is estimated from the difference between the distance from the peak center to the left base (L) and the distance from the peak center to the right base (R) as follows:

Skewness, $K = \frac{|L-R|}{L+R+\epsilon}$

Lower K values indicate more symmetric peaks, while higher values indicate increased asymmetry. To robustly compare the skewness of all detected peaks within the distribution, a robust z-score ($Z_k$) is computed using the median and Median Absolute Deviation (MAD) of all skewness values:

$Z_k = \frac{K - median(K)}{MAD(K)}$

where:

MAD(K) = $median(|K - median(K)|) + \epsilon$

Finally, the skewness-based quality weight ($Q_K$) is obtained by applying a sigmoid transformation to the robust z-score:

$$Q_K = \frac{1}{1 + e^{Zk}}$$

This transformation softly rewards symmetric peaks ($Z_k < 0$) with higher weights and penalizes highly asymmetric peaks ($Z_k > 0$) without introducing hard thresholds.

This score is calculated only for peaks contributing at least 2% of the total area under the distribution. The top two peaks, ranked by their score values, are then evaluated using additional height and area thresholds. Both peaks are required to contribute at least 5% of the total distribution area, or alternatively, the smaller peak is required to have a height of at least 15% of the height of the major peak.

If neither of these conditions is satisfied, the minor peak is further evaluated based on its position relative to the major peak along the X-axis, representing the copy number. Specifically, the minor peak is required to occur on the right side of the major peak, indicating a potentially expanded or longer allele, thereby improving sensitivity for detecting repeat expansions. If this condition also fails, the remaining cluster is re-examined for the presence of subclusters using HDBSCAN clustering, with the minimum cluster size set to either 10 reads or 20% of the total supporting reads, whichever is greater. If multiple subclusters are identified, the two largest clusters are selected as the representative alleles. Otherwise, the locus is considered homozygous, and only a single allele is reported in the VCF.

#### Consensus sequence generation

After clustering the reads, the repeat sequences from each cluster underwent partial order alignment using the abPOA (Gao et al., 2021) tool to generate a consensus sequence. Because graph construction is performed iteratively, the order in which sequences are incorporated into the graph could influence the resulting consensus sequence.

In WGS mode, sequences are arranged in descending order based on their frequency prior to consensus generation. In Amplicon mode, sequences are ranged alternately from both sides of the median sequence length to reduce length-based bias during graph construction.

If the median allele sequence length exceeds 10 kb, a position-wise consensus approach is used instead, where the most frequent base at each position across all supporting reads is assigned as the final consensus sequence.

#### VCF output

After phasing the reads supporting a given locus, locus-level statistics are compiled and recorded in the VCF file. These include the genotype alleles, repeat copy number for each allele, the longest pure repeat motif and its copy number, phased read depths, the central 95% percentile range of allele lengths within each haplogroup, mean methylation levels for each allele, and the number and base qualities of SNVs used for phasing.

#### Data pruning for memory-optimisation

ATaRVa implements several memory-management strategies to retain only the data required for downstream processing, thereby minimizing memory usage and improving computational efficiency during large-scale repeat genotyping. Once a repeat locus has been genotyped, all locus-specific information is removed from memory. Similarly, if the start position of the current read exceeds the end position of a previously stored read and that read does not enclose any ungenotyped repeat loci, the read and its associated data are discarded. Any SNVs supported by the removed read have their coverage counts updated accordingly within the SNV data structure, and SNV positions whose coverage decreases to zero are subsequently removed.

#### Merging VCFs

For population-scale and multi-sample analyses, individual VCF files must be merged such that sample genotypes are updated relative to the complete set of alleles observed across all samples. Additionally, population-level annotations, including AN (the number of genotyped alleles at a locus) and AC (the allele count for each alternate allele across all samples), must be recalculated along with other INFO fields. In the merged VCF, a separate sample column is added for each individual following the nine mandatory VCF columns. ATaRVa provides a dedicated merging module designed to efficiently handle large cohorts comprising thousands of VCF files and millions of loci (Fig S10). This module requires BGZF-compressed and indexed VCF files together with the BED file defining the regions of interest.

The merging process begins by partitioning the VCF records into equally sized genomic chunks and distributing the input VCFs across multiple processors. Sample names are extracted from each VCF, and the records are loaded into Polars DataFrames for efficient processing. Based on the order of the input files, the first VCF serves as the reference framework, storing common locus-level information such as chromosome, start and end coordinates, reference repeat sequence, and repeat motif. For subsequent VCFs, only sample-specific information, including alternate allele sequences, read support, methylation metrics, and motif decomposition information, is extracted and aligned with the existing DataFrame using the genomic coordinates as merge keys.

The available computational threads are divided between reader and processing tasks such that each reader process is assigned a similar number of VCF-processing threads. The BED file is partitioned into genomic intervals, and each reader process is assigned a subset of tandem repeat loci. Intermediate Parquet files are generated whenever data for approximately 5,000 loci across 200 samples have been accumulated. Once all samples corresponding to a given set of loci have been read, the intermediate Parquet files are merged into a single DataFrame.

The merged DataFrame is then distributed across multiple processing threads to generate the final multi-sample VCF records. During this step, sample genotypes are recalculated relative to the complete set of alternate alleles observed at each locus. Newly observed alleles are incorporated into the ALT field, genotypes are updated accordingly for all samples, and population-level statistics such as AN and AC are recomputed. The processed records are subsequently written as intermediate VCF files. After all processing threads associated with a reader process have completed, their intermediate VCF files are merged vertically to produce a reader-level VCF. Finally, the reader-level VCFs from all genomic partitions are concatenated to generate the final merged multi-sample VCF file.

### VisuaMiTRa workflow

#### Input Parsing and Query Filtering

VisuaMiTRa processes tabix-indexed Variant Call Format (VCF) files alongside their corresponding index (.tbi) files. The workflow initiates when a user queries for specific samples and/or filters for a target genomic window by optionally specifying a chromosome, start & end coordinates. The backend layer (**Python** & **FastAPI**) establishes a file handle over the indexed VCF. It executes validation checks, intercepting queries for empty regions before data transmission. To stream large, multi-sample datasets without excessive memory consumption, the system uses a token-based pagination mechanism. The backend encodes the genomic position of the last row processed in a given page, into a Base64 cursor, within the HTTP response headers. For subsequent data requests, this cursor is decoded to resume streaming exactly where the previous page halted. If the query range spans across chromosome boundaries, a natural sorting algorithm increments the file pointer to the next consecutive contig, maintaining an uninterrupted data stream across the genome.

#### Repeat Sequence Decomposition & Methylation

The ATaRVa pipeline handles the underlying sequence decomposition and methylation to generate structured formatting tags for the frontend layer. The frontend parses the bifurcated components of **DS** tag to extract structural copy numbers, repeat variations, and exact sequence coordinate sub-arrays. These metrics are dynamically mapped onto a shared horizontal grid axis to render proportional, color-coded block layouts, enabling instantaneous cross-cohort comparison of motif expansions and locus shifts. The **MV** tag delivers Base64-encoded strings containing both the relative CpG positions and their methylation values, which the backend decodes into coordinate-percentage matrices (0% to 100%), so the frontend can loop through them, plotting every CpG site along the allele through a continuous color gradient palette.

#### User Interface

Along with the decomposition & methylation information, the backend derives metadata of the selected samples from the vcf headers. Metadata includes sample names, genotypes, read-support & mean-methylation. All the acquired data is streamed as a tab-separated value (TSV) matrix to the front-end user interface and the layout engine mounts data as synchronized **React** components. VisuaMiTRa allows simultaneous comparison of multiple samples, using a searchable dropdown and a synchronized genomic scale. Users can navigate the genome via a precise coordinate location picker paired with a chromosome ideogram that highlights the exact physical cytogenetic location of the active target repeat. A collapsible dashboard displays sample metadata in a compact, three-row layout that expands dynamically on demand. The main plotting area features interactive scale controls and toggleable tabs to seamlessly switch between sequence decomposition and methylation profiles, automatically scaling all tracks relative to the longest allele in the batch. For publication-ready outputs, the interface provides comprehensive visual customization—including motif colors, hover tooltips, font adjustments, and methylation color scales (like viridis), alongside a high-resolution export engine supporting vector (SVG) and raster (PNG/JPEG) downloads with optional metadata inclusion.
